# CRISPR-Cas9-mediated knockout of *CYP79D1* and *CYP79D2* in cassava attenuates toxic cyanogen production

**DOI:** 10.1101/2021.10.08.462827

**Authors:** Michael A. Gomez, Kodiak C. Berkoff, Baljeet K. Gill, Anthony T. Iavarone, Samantha E. Lieberman, Jessica M. Ma, Alex Schultink, Stacia K. Wyman, Raj Deepika Chauhan, Nigel J. Taylor, Brian J. Staskawicz, Myeong-Je Cho, Daniel S. Rokhsar, Jessica B. Lyons

## Abstract

Cassava (*Manihot esculenta* Crantz) is a starchy root crop that supports over a billion people in tropical and subtropical regions of the world. This staple, however, produces toxic cyanogenic compounds and requires processing for safe consumption. Excessive consumption of insufficiently processed cassava, in combination with protein-poor diets, can have neurodegenerative impacts. Reducing the cyanogen content by conventional breeding is problematic due to the heterozygous nature of the crop; recombination will generally disrupt a clonally propagated cultivar’s suite of desirable traits. To reduce cyanide levels in cassava, we used CRISPR-mediated mutagenesis to disrupt the cytochrome P_450_ genes *CYP79D1* and *CYP79D2* whose protein products catalyze the first step in cyanogenic glucoside biosynthesis. Knockout of both genes eliminated cyanide in leaves and storage roots of cassava accession 60444 and the West African, farmer-preferred cultivar TME 419. Although knockout of *CYP79D2* alone resulted in significant reduction of cyanide, mutagenesis of *CYP79D1* did not, indicating these paralogs have diverged in their function. Our work demonstrates cassava genome editing for food safety, reduced processing requirements, and environmental benefits that could be readily extended to other farmer-preferred cultivars.

## Introduction

The starchy root crop cassava (*Manihot esculenta* Crantz, also known as tapioca, yuca, or manioc, among others) is an important staple for over a billion people in tropical and subtropical regions of the world, including roughly 40% of Africans (Lebot, 2019; Nweke, 2004). It is an excellent food security crop due to its tolerance for drought and marginal soils, and because its tuberous roots can be left in the ground and harvested when needed (Howeler *et al*., 2013). A major challenge, however, is the presence of toxic cyanogenic compounds (e.g., cyanogenic glucosides) in cassava, which must be removed by post-harvest processing to prevent cyanide exposure. The processing of roots can be laborious, and in Africa falls disproportionately on women and girls (Chiwona-Karltun *et al*., 1998; Curran *et al*., 2009; Ogunwande *et al*., 2016). Processing of cassava roots and leaves also results in nutrient loss, thereby reducing the food value of processed cassava products (Boakye Peprah *et al*., 2020; Hawashi *et al*., 2019; Maziya-Dixon *et al*., 2009; Montagnac *et al*., 2009).

Cyanogenic glucosides are broken down to release the toxin cyanide following cellular disruption (e.g., during digestion). Distributed throughout the body via the bloodstream, cyanide halts mitochondrial electron transport, thereby preventing cells from using oxygen to produce energy and causing cell death (Dobbs, 2009). The central nervous system is particularly impacted by this toxin due to its substantial oxygen demand. The risks of insufficient cassava processing include acute cyanide poisoning which can be fatal. Chronic cyanide exposure from dietary intake induces the paralytic disease konzo, is associated with neurodevelopmental deficits, and exacerbates tropical ataxic neuropathy and goiter (Kashala-Abotnes *et al*., 2018; Nhassico *et al*., 2008; Nzwalo and Cliff, 2011; Tshala-Katumbay *et al*., 2016). Sulfur-containing amino acids are required to detoxify cyanide in the body; thus, those with a protein-poor diet heavily reliant on cassava are particularly at risk for adverse effects from cyanide exposure (Nzwalo and Cliff, 2011). Konzo is more likely to occur in women of childbearing age and children (Baguma *et al*., 2021).

Processing to remove cyanogenic content from tuberous roots can be achieved by chipping and air drying, grinding, mashing and steeping, and/or fermentation. All require 24 hours to several days to complete, with premature consumption exposing consumers to risk. Insufficient processing may be driven by lack of access to water, hunger, and other factors such as drought and social unrest. Industrial scale processing of cassava poses risks to the environment and to workers through cyanide release into wastewater and the air, respectively (Adewoye *et al*., 2005; Dhas *et al*., 2011; Ehiagbonare *et al*., 2009). Cyanide levels above WHO recommendations have been found in commercial cassava products as well as household flour (Burns *et al*., 2012; Kashala-Abotnes *et al*., 2018).

Cassava was domesticated in the southern Amazon Basin over 8,000 years ago (Clement *et al*., 2010; Olsen and Schaal, 1999; Pearsall, 1992; Watling *et al*., 2018) and has been grown in Sub Saharan Africa for fewer than 500 years (Hillocks, 2002; O’Connor, 2013). Processing approaches vary by region and specific cultivated variety (cultivar) used. There are cultural preferences for growing high cyanogenic cultivars (known as “bitter” cultivars) in some contexts, for example to deter theft (Chiwona-Karltun *et al*., 1998). Shortcuts are sometimes taken during processing, especially when food is in short supply (Banea *et al*., 1992; Essers *et al*., 1992; Fitzpatrick *et al*., 2021; McKey *et al*., 2010). A mismatch between expected and actual cyanide levels (due to use of a different cultivar or environmental factors) may render the usual processing insufficient. Notably, levels of cyanogenic molecules (hereafter cyanogens) are higher under drought stress (Brown *et al*., 2016; El-Sharkawy, 1993; Okogbenin *et al*., 2003; Vandegeer *et al*., 2013). Droughts are becoming more frequent and severe (Kendon *et al*., 2019), thus putting cassava consumers at added risk of cyanide exposure.

The pathway for synthesizing cyanogenic glucosides requires cytochrome p450 (CYP) enzymes of the CYP79 family (Luck *et al*., 2016). CYP79D proteins catalyze the first, limiting step of synthesis for cassava’s principal cyanogens, linamarin and lotaustralin (Andersen *et al*., 2000). These are synthesized in leaves and shoots and then transported into the storage roots (Jørgensen *et al*., 2005) (**Figure 1a**). CYP79D is encoded in cassava by the paralogous genes *CYP79D1* and *CYP79D2* (Andersen *et al*., 2000), which arose through the whole-genome duplication found in this lineage (Bredeson *et al*., 2016). Linamarin (derived from valine) accounts for greater than 80% of cassava cyanogens (Cereda and Mattos, 1996).

**Figure 1.**
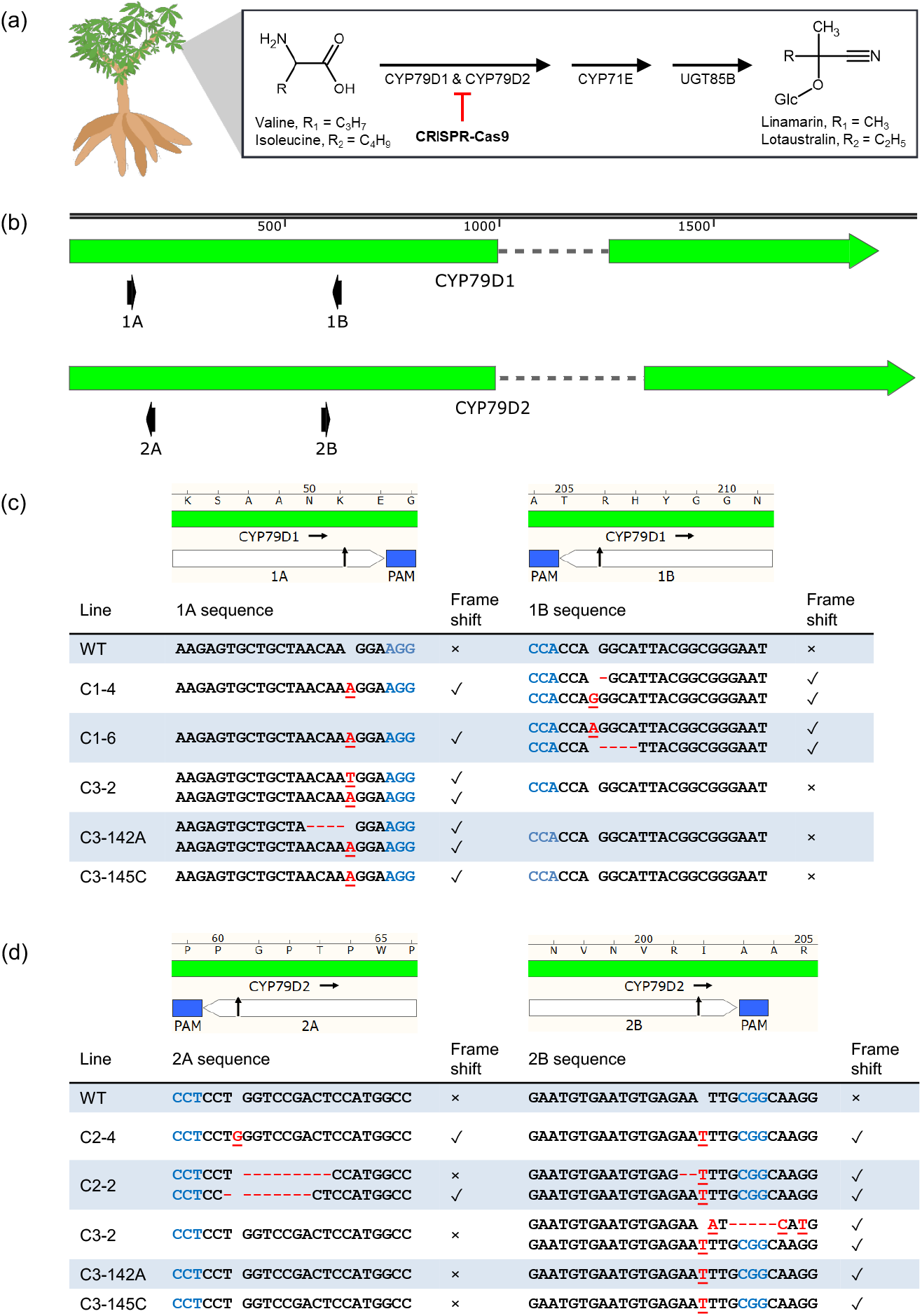
CRISPR-Cas9 induces indels at *CYP79D1* and *CYP79D2* gRNA target sites in transgenic 60444 lines. Cassava biosynthetic pathway for the cyanogenic glucosides linamarin and lotaustralin. This process primarily occurs in the leaves. The step catalyzed by CYP79D1 and CYP79D2 enzymes was selected for disruption by the CRISPR-Cas9 system. Respective side chains are labeled as R1 and R2. Glc, glucose. Chemical structures created using ACD/Chemsketch (ACD/Chemsketch, version 2021.1.1, 2021). Lengths of *CYP79D1* and *CYP79D2* genes are to nucleotide scale (top bar). Exons are denoted by solid blocks and introns are represented as dashed lines. Arrowheads indicate the 3′ terminus. Diagrams of the protospacers (white) and protospacer adjacent motifs (PAMs, blue) of *CYP79D1* (c) and *CYP79D2* (d) gRNA targets are aligned to edited line genotypes. Edited lines are identified by the CRISPR construct with which they were modified (C1, C2, C3), followed by an index number (e.g. 142A). Black arrow indicates predicted CRISPR-Cas9 cut site. Lengths are to amino acid (top bar) and nucleotide (bottom table) scale. Homozygous genotypes are shown as a single sequence per line. Bi-allelic genotypes are shown as two sequences per line. Mutations on the same haplotype at 1A and 1B sites in a given mutant line are shown in the same row. Insertions are denoted by red, underlined nucleotides. Deletions are denoted by red dashes. Presence of a frameshift mutation at the corresponding target site is denoted by ✓; absence of a frameshift mutation is denoted by ×. WT, wildtype. Maps created with SnapGene.

Cyanogens play multiple roles in plants including defense and metabolism. Although cyanogens can deter herbivores (Bernays *et al*., 1977; Gleadow and Møller, 2014; Rajamma and Premkumar, 1994), this is not the case for cassava in all contexts or against all herbivores, possibly due to coevolution (Pinto-Zevallos *et al*., 2016; Riis *et al*., 2003). For example, the whitefly *Bemisia tabaci* detoxifies cyanogenic glucosides by enzymatic conversion to inert derivatives (Easson *et al*., 2021). It has been proposed that cyanogens shuttle reduced nitrogen to cassava roots for protein synthesis; increased nitrate reductase activity in roots, however, may compensate for reduced cyanogen availability (Jørgensen *et al*., 2005; Narayanan *et al*., 2011; Siritunga and Sayre, 2004; Zidenga *et al*., 2017). Cyanogens are also hypothesized to play a role in initiating postharvest physiological deterioration of the roots by triggering reactive oxygen species production (Zidenga *et al*., 2012). Modulation of cyanide levels may, therefore, bolster the longevity of harvested roots. Generation of acyanogenic cassava will facilitate further investigation of cyanogenic potential in these roles.

Cyanogen production varies naturally among cultivars (see, *e*.*g*., (Ogbonna *et al*., 2021; Ospina *et al*., 2021; Whankaew *et al*., 2011)), indicating that cyanogen levels can be modulated without disrupting other desirable plant properties. Knockdown of the *CYP79D* genes using RNAi has been shown to reduce cyanogen levels in cassava leaves and roots (Jørgensen *et al*., 2005; Piero, 2015; Siritunga and Sayre, 2003). Knockdown plants displayed a wildtype phenotype when grown in soil (Jørgensen *et al*., 2005; Piero, 2015; Siritunga and Sayre, 2003). Taken together, these observations indicate that cyanogen levels can be modulated without disrupting other desirable plant properties.

Here, we show that the toxic cyanogenic glucosides in cassava can be eliminated by knocking out the *CYP79D* genes. We used CRISPR-Cas9 to edit the CYP79 genes, singly and in combination, in the model variety 60444 and the popular West African landrace TME 419. *Agrobacterium*-mediated CRISPR-Cas9 editing is efficient in cassava and has been demonstrated in several cultivars (Bull *et al*., 2018; Gomez *et al*., 2019; Hummel *et al*., 2018; Odipio *et al*., 2017; Veley *et al*., 2021). We show the reduction of cyanogen levels through targeted knockout of one or both of the *CYP79D* genes. Our targeted genome editing approach provides a complete loss of function without altering genes related to other desirable traits; this is important in a vegetatively propagated crop like cassava whose improvement by conventional breeding is laborious. We find that dual knockouts eliminate cyanogenic potential in both cassava accessions. Single gene knockout lines reveal differential contribution of the two *CYP79D* genes to cassava cyanogenesis. The knockout lines described here will facilitate further research into the cyanogenesis pathway, and provide experimental materials for controlled field studies on the role of cyanogens in cassava.

## Results and Discussion

### Site-specific mutation of *CYP79D* genes by transgenic expression of sgRNA-guided Cas9

To disable CYP79D activity we designed three gRNA constructs to target *CYP79D1* and *CYP79D2*, singly and simultaneously, using CRISPR-Cas9 editing (**Table 1, Figure S2, Experimental Procedures**). We targeted two sites per gene to induce gene disruption by frameshift mutation at either site and/or excision of a substantial portion of the gene (**Figure 1b**). Multiplex editing capability of the CRISPR-Cas9 system was boosted by employing the tRNA-processing system for gRNA expression (Xie *et al*., 2015).

**Table 1.** CRISPR constructs used in this work.

| **CRISPR Construct** | **Target Gene(s)** | **gRNAs** |
| --- | --- | --- |
| C1 | *CYP79D1* | 1A, 1B |
| C2 | *CYP79D2* | 2A, 2B |
| C3 | *CYP79D1*, *CYP79D2* | 1A, 1B, 2A, 2B |

We confirmed *in planta* functionality of the CRISPR constructs by using the surrogate model *Nicotiana benthamiana*. For sensitive detection of Cas9 activity, we adapted a geminivirus system (Baltes *et al*., 2014) to amplify target sequences in the plant cell, thereby facilitating generation of many edited target sequences following cleavage by Cas9. Target regions of *CYP79D1* and *CYP79D2* were cloned into a plasmid encoding a geminivirus replicon (**Figure S1a**). Following *Agrobacterium*-mediated co-delivery and expression of these gemini-vectors and our CRISPR constructs in leaves (**Figure S1b**), simultaneous cleavage and excision of the DNA between the two target sites by active Cas9 and gRNAs resulted in a shorter target region amplicons (**Figure S1c,d**), confirming that the CRISPR constructs were functional.

To generate transgenic cassava lines with disrupted CYP79D genes, we transformed friable embryogenic calli (FEC) from cassava accessions 60444 and TME 419, using *Agrobacterium* carrying our confirmed constructs. For each construct, we recovered multiple independent T0 transgenic plant lines. We characterized the CRISPR-Cas9 induced mutations in these lines using Sanger sequencing of the target regions (**Figure 1b, Figure 1c, Figure S3, File S1**). Mutagenesis of targets 1A and 2B occurred at a higher frequency than of 1B and 2A, which may be due to differences in the gRNA sequences and their respective binding efficiencies. Rarely was the region between the two target sites within a gene deleted. This result was unexpected since this excision, by all CRISPR constructs, was easily detectable in the *N. benthamiana* surrogate assay. The smaller size of the excision product may have led to preferential PCR amplification in this assay.

We sorted edited lines into the following mutation categories: bi-allelic (carrying two different mutations, one for each copy of the targeted gene); homozygous (having two identical mutations of their alleles); and heterozygous (carrying one mutagenized allele and one wildtype allele). Plants that carry more than two sequence patterns, indicating chimerism, were designated “complex.” Plants with complex genotypes may still yield functional transcripts and therefore pose a challenge to phenotype prediction. We selected mutant lines showing bi-allelic or homozygous frameshift mutations leading to premature stop codons for further analysis (**Figure SAAL**). To confirm the genotypes from selected lines, we sequenced the target regions using Illumina sequencing (**Experimental Procedures**). TME 419 line C3-10 appeared from Sanger sequencing to be a double knockout mutant; the amplicon sequencing data, however, showed 6.9% wildtype reads at site 2B, revealing that this line is mosaic, with some cells having an intact *CYP79D2* gene (**Figure S7**). We found no evidence of off-target mutagenesis (**Table S3**) in 60444-derived edited lines based on the sequencing of potential off-target sites for gRNA targets 1A, 1B, 2A, and 2B (**Experimental Procedures, Table S1**). Sequencing of cDNA from targeted *CYP79D1* and *CYP79D2* loci in 60444-derived lines showed that their RNA transcripts had the expected sequences.

### Cyanogenic potential of edited plants

#### LC-MS cyanogen measurements from *in vitro* plantlets

We measured levels of linamarin and lotaustralin in leaves of edited 60444 and TME 419 *in vitro* plantlets using liquid chromatography-mass spectrometry (LC-MS), with age-matched wildtype *in vitro* plantlets as positive controls (**Experimental Procedures, Figure S5, Figure S6**). A single leaf sample was taken from one to three plants per line, and an average value for each plant calculated from three technical replicates. No linamarin or lotaustralin was detected in negative (no tissue) controls.

Linamarin was not detected in dual knockout lines derived from either 60444 or TME 419 (**Figure S5a, Figure S6a**). Linamarin values of a 60444-derived *CYP79D1* knockout line (0.12–0.17 g per kg fresh weight [g/kgfw]) fell within the range of wildtype values (0.11–0.40 g/kgfw). Linamarin values for three TME419-derived *CYP79D1* knockout lines were more variable, ranging from undetectable to 0.47 g/kgfw, while wildtype ranged from 0.15–0.23 g/kgfw. *CYP79D2* knockouts, however, in both accessions displayed linamarin levels consistently much lower than wildtype: highest at 0.03 g/kgfw in 60444 and 0.10 in TME419, with two out of three TME419 lines at or below 0.01 g/kgfw. A low level of linamarin was detected in TME 419 line C3-10 (0.003–0.020 g/kgfw), consistent with the mosaicism for a wildtype *CYP79D2* allele shown by Illumina sequencing. We did not consider this line further.

Lotaustralin measurements from *in vitro* plantlets generally followed the trend seen for linamarin, except at much lower levels (**Figure S5a, Figure S6a**). Wildtype values reached 0.019 g/kgfw in 60444 and 0.007 g/kgfw in TME419. Low lotaustralin levels were expected, since linamarin is the predominant cyanogen in cassava (Nartey, 1968). Lotaustralin was only detected in one *CYP79D2* knockout plant of one line in each of the two accessions. Extremely small amounts (0.0005 and 0.0007 g/kgfw) of lotaustralin were detected in two 60444 double knockout plants. As no linamarin was detected in any sample from these lines, and no lotaustralin was detected in the other plants from these lines, it is unlikely that these readings reflect true cyanogenesis but may instead be the result of technical contamination.

#### Cyanide measurements from adult plants

In addition to plantlets, we also measured cyanide levels in leaves and tuberous roots of wildtype and mutant 60444 and TME 419 adult plants, using a picrate assay (Bradbury *et al*., 1999) (**Figures 2 and 3**). For each accession, plants were grown in synchronous cohorts, transferred to soil, and grown in a glasshouse. Assays were performed 6–8 months after soil transfer, with a whole cohort of the same accession removed from the greenhouse and assayed on the same day. Edited plants were morphologically indistinguishable from wildtype (**Figure S4**). Cyanide content can vary considerably between roots of the same plant and plants of the same cultivar (Cooke *et al*., 1978); in expectation of this variability, up to nine root samples from at least three plants per line were analyzed. As seen in *in vitro* plantlets, dual knockout lines had no cyanogenic potential, while *CYP79D2* knockouts had a more drastic reduction in cyanogenic potential relative to wildtype than did knocking out *CYP79D1* knockouts (**Figure 2a,b; Figure 3a,b**).

**Figure 2.**
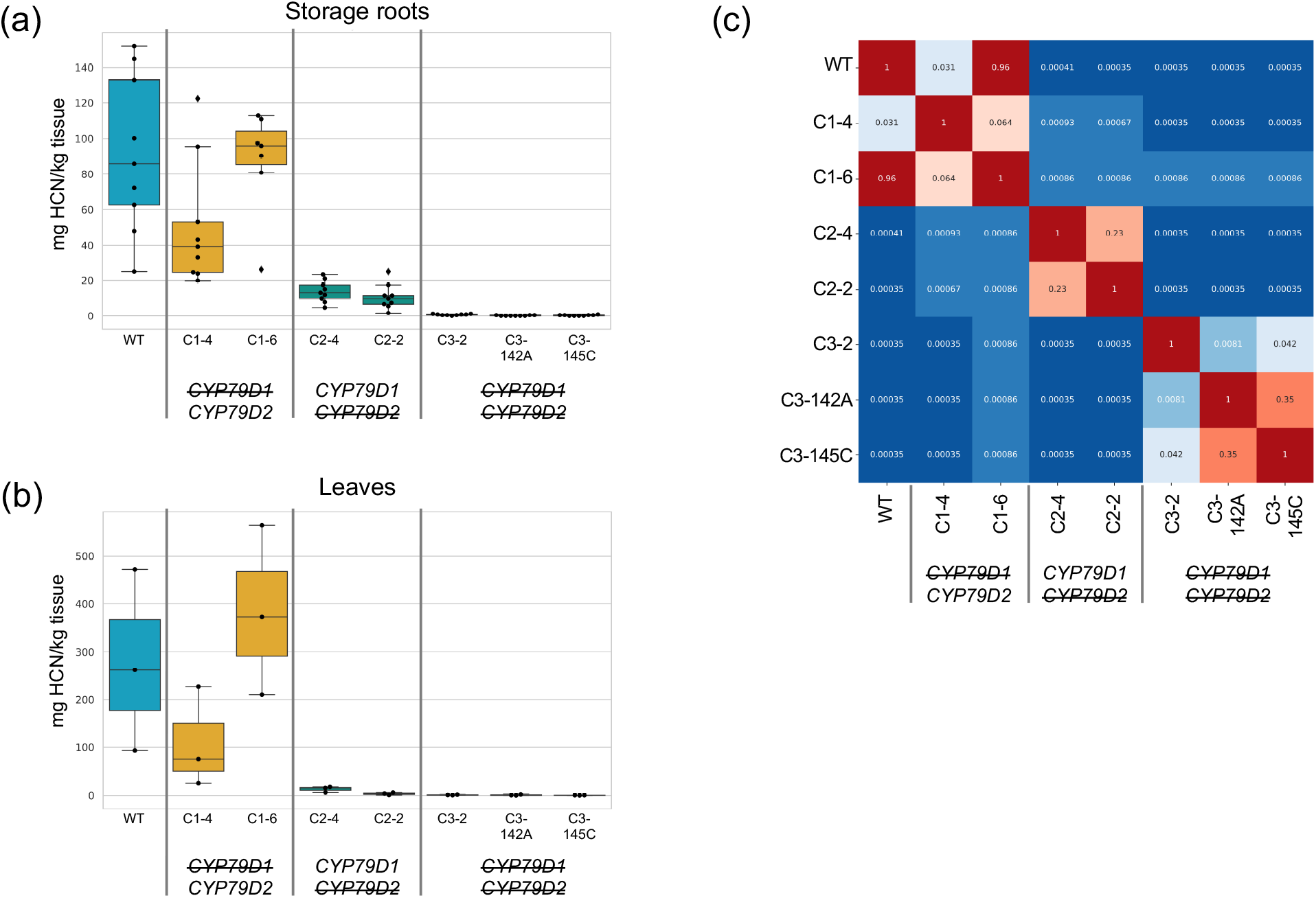
Dual knockout of *CYP79D1* and *CYP79D2* eliminates cyanide production in accession 60444. Assays were conducted 6.5 months after plants were transferred to soil. (a and b) Box and whisker plots of cyanide values in 60444 tuberous roots (a) and leaves (b) in mg HCN per kg tissue, as detected by picrate assay. Black dots are biological replicates. The median, and lower (25th percentile) and upper (75th percentile) quartiles are indicated. Whiskers define the minimum and maximum regions of the data; data points outside of these are outliers. (c) Wilcoxon Rank Sum p-values calculated from pairwise comparisons between lines of root cyanide content. Values less than 0.05, indicating the distributions of the values are statistically different between the two lines, are colored in shades of blue. Values greater than 0.05 are colored in shades of red. WT, wildtype. Edited lines are identified by the CRISPR construct with which they were modified (C1, C2, C3), followed by an index number (e.g. 142A).

**Figure 3.**
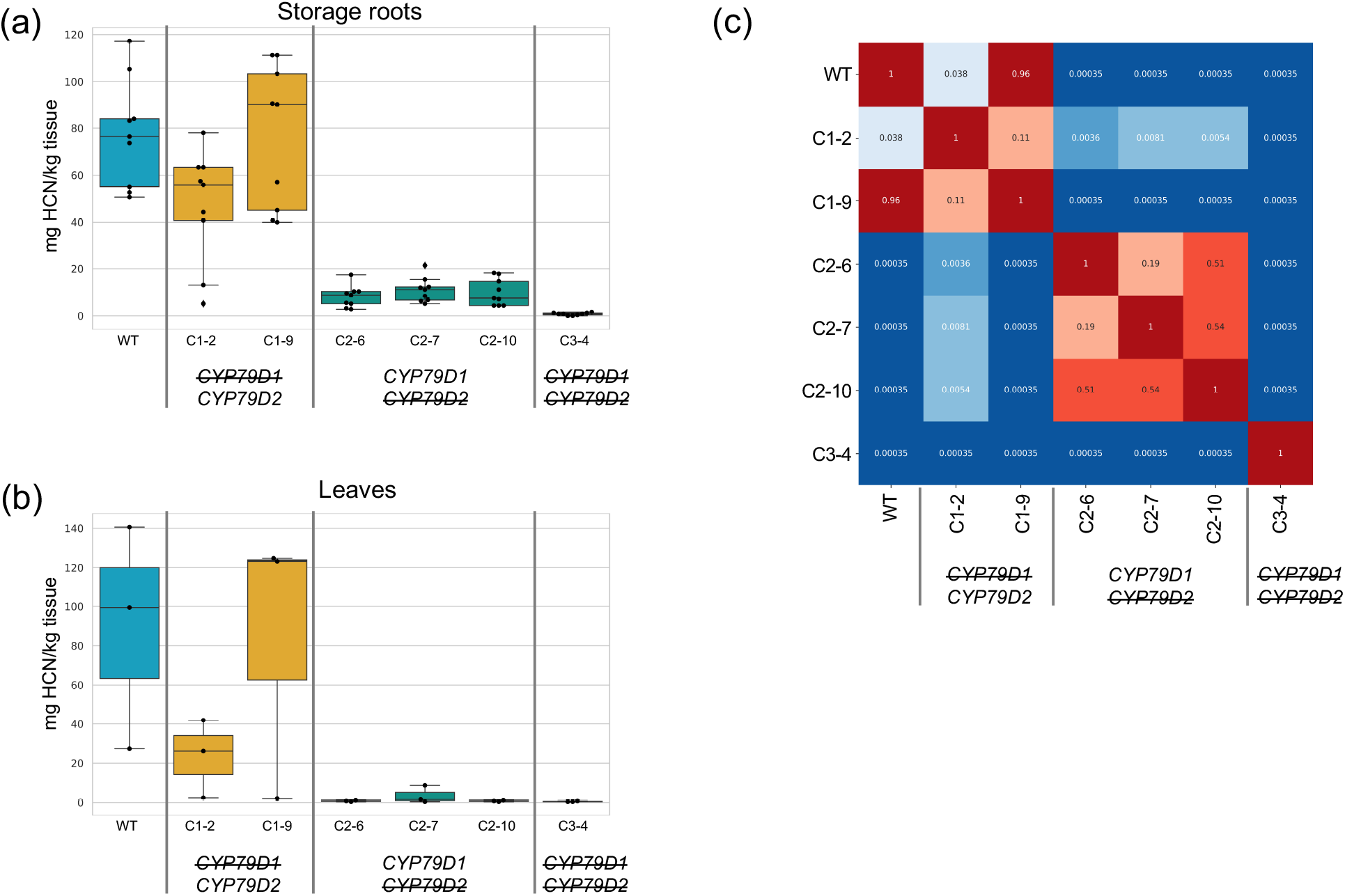
Dual knockout of *CYP79D1* and *CYP79D2* eliminates cyanide production in accession TME 419. Assays were conducted 8 months after plants were transferred to soil. (a and b) Box and whisker plots of cyanide values in TME 419 tuberous roots (a) and leaves (b) in mg/kg tissue, as detected by picrate assay. Black dots are biological replicates. The median, and lower (25th percentile) and upper (75th percentile) quartiles are indicated. Whiskers define the minimum and maximum regions of the data; data points outside of these are outliers. (c) Wilcoxon Rank Sum p-values calculated from pairwise comparisons between lines of root cyanide content. Values less than 0.05, indicating the distributions of the values are statistically different between the two lines, are colored in shades of blue. Values greater than 0.05 are colored in shades of red. WT, wildtype. Edited lines are identified by the CRISPR construct with which they were modified (C1, C2, C3), followed by an index number.

The observed variability within a given line was expected, as noted above. Hence, we performed Wilcoxon Rank Sum tests on the root data, using pairwise comparisons to test whether cyanide levels in different lines are distinct from one another (**Figure 2c, Figure 3c**). In dual knockout lines generated from 60444 and TME 419, the zero (or very near zero) assayed cyanide levels were distinct from those of wildtype as well as those of all single knockout lines. We had two *CYP79D1* knockout lines from each accession. In both accessions, one of the *CYP79D1* knockout lines had cyanide levels distinct from wildtype and the other did not. This is consistent with our observation that knocking out *CYP79D1* alone does not reliably reduce cyanogenic potential below wildtype levels. In both accessions, all *CYP79D2* knockout lines had cyanide levels distinct from wildtype, *CYP79D1* knockouts, and dual knockouts. This is consistent with our assessment that knocking out *CYP79D2* alone provides a near complete, but not total, reduction in cyanogenic potential.

The difference in reduction of cyanide resulting from single knockouts of *CYP79D1* and *CYP79D2* indicates a disparity in their relative contributions to cassava cyanogenesis. Differences in gene expression and/or protein sequence may explain this disparity. *CYP79D2* shows higher transcriptional activity than *CYP79D1* (Wilson *et al*., 2017) (**Figure S8**). The 1000-bp regions upstream of the *CYP79D1* and *CYP79D2* transcript sequences have 50.17% identity; at the amino acid sequence level, CYP79D1 and CYP79D2 have 85.74% identity (Bredeson *et al*., 2021). Multiple amino acid mismatches lie in the transmembrane and p450 domains predicted by SMART (Letunic *et al*., 2021). It is also possible that genetic and metabolic feedback loops are influencing cyanogenic output of the intact *CYP79D* gene in single knockout lines. Gene and protein expression and regulation of the *CYP79D* genes merit further study.

Cyanide levels are highly variable across accessions (Ogbonna *et al*., 2021). To identify farmer-preferred cultivars that could be future targets for *CYP79D* gene modification, we measured leaf and root cyanide content in accessions 60444, TME 419, TMS 91/02324, TMS 98/0505, TME 3, MCol 22, and MCol 2215 (**Figure 4**). These all have established transformation protocols (Chauhan *et al*., 2018; Li *et al*., 1996; Siritunga and Sayre, 2003; Taylor *et al*., 2012; Zainuddin *et al*., 2012). Ranges of storage root cyanide values of the different accessions largely overlapped. Relative cyanide content in leaves was not predictive of relative cyanide content in roots across accessions. This result is consistent with previous work showing weak correlation between leaf and storage root cyanide content (Ospina *et al*., 2021). Therefore, leaf cyanide levels should not be used to determine relative root levels between accessions. The observed variability indicates that there are differences between cassava accessions in terms of cyanogen biosynthesis, transport and/or metabolism. Understanding the mechanism and regulation of these differences could be useful for modulation of this pathway in the future.

**Figure 4.**
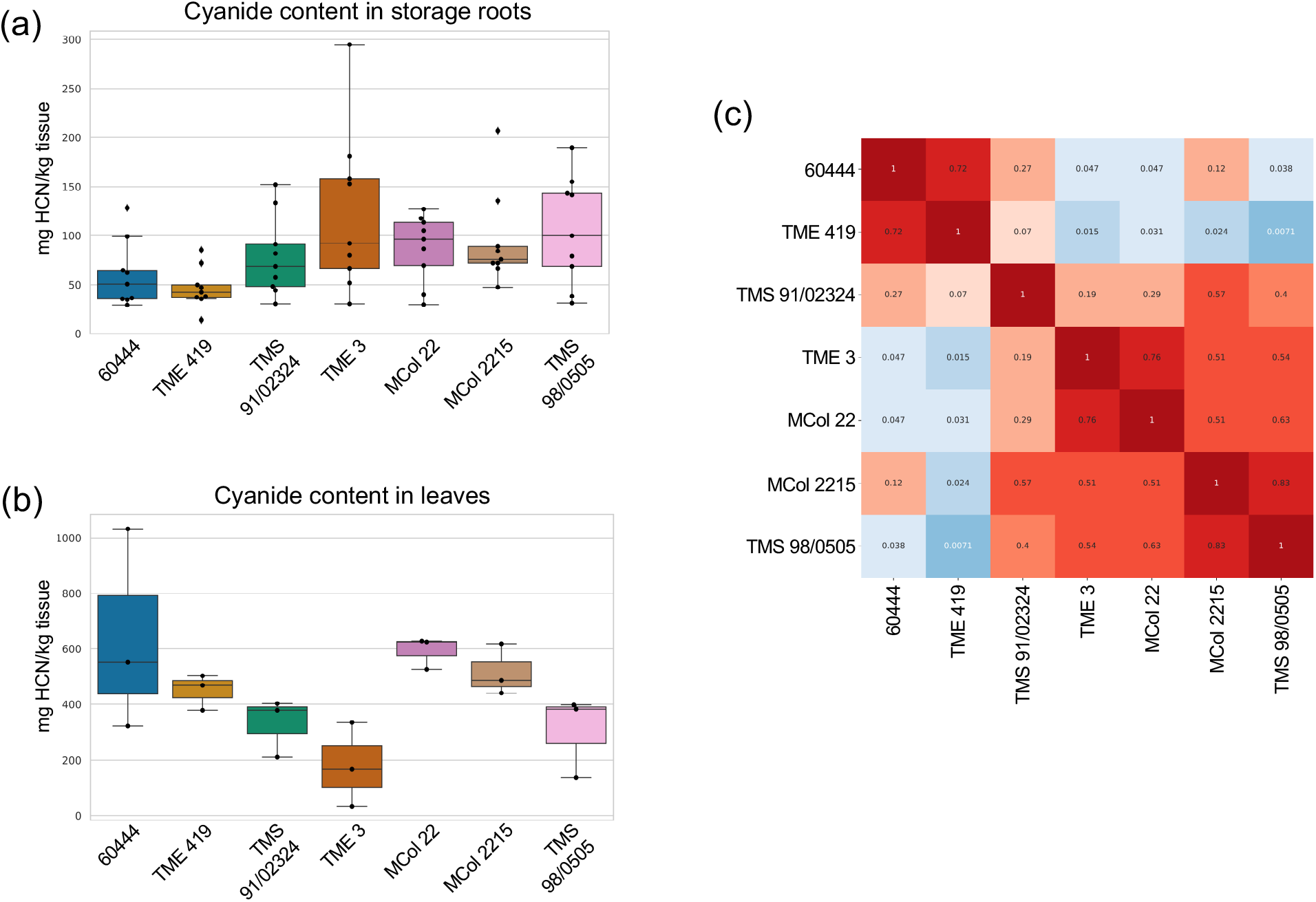
Cyanide levels from seven transformable cassava accessions. Assays were conducted seven months after plants were transferred to soil. (a and b) Cyanide levels in (a) storage roots and (b) leaves of the indicated cassava accessions in mg HCN per kg tissue, as detected by picrate assay. Black dots are assayed values. The median, upper, and lower quartiles are indicated. Whiskers define the minimum and maximum regions, and dots outside of these are outliers. (a) Root measurements. For each accession, samples were taken from a total of nine tuberous roots, from four to six plants. (b) Cyanide measured from leaf samples. For each accession, a total of three leaf samples were taken, from three plants. (c) Wilcoxon Rank Sum p-values calculated from pairwise comparisons between accessions of root cyanide content. Values less than 0.05, indicating the distributions of the values are statistically different between the two lines, are colored in shades of blue. Values greater than 0.05 are colored in shades of red.

## Conclusions

Genome editing is a powerful and heritable way to disable genes of interest for functional assessment and crop improvement. This study marks the first report of engineering acyanogenic plants via the CRISPR-Cas system. Here, we targeted cyanogenesis genes *CYP79D1* and *CYP79D2* to achieve reduction in cassava’s cyanogenic potential. We demonstrated a loss of cyanogenic potential in four dual knockout lines derived from two cassava accessions, via measurement of cyanogenic glucosides in *in vitro* plantlets as well as evolved cyanide in adult leaves and storage roots.

Cassava is paleotetraploid, and has two genes that encode the enzyme CYP79D (Andersen *et al*., 2000; Bredeson *et al*., 2016). Gene duplication provides an evolutionary substrate for functional diversification (Otto and Yong, 2002), since each gene can then accumulate novel and/or complementary mutations, including variation in substrate specificity and gene expression. The replacement of a single gene by a group of paralogs opens the door to subtle regulation of biochemical networks. We tested this in cassava using the *CYP79D* genes, and our data suggest that *CYP79D2* is responsible for a greater proportion of CYP79D enzymatic function than *CYP79D1*. It will be interesting to see whether this is due to differences in expression at the RNA or protein level, or in the efficacy of the protein itself. There may also be differences in spatial and/or temporal expression between these genes. *CYP79D2* knockouts showed a seven-to ten-fold reduction of cyanide content, when comparing the mean values of these edited lines to mean values from the corresponding wildtype accessions. Thus, knockout of *CYP79D2* alone provides a straightforward mechanism for generating stably low-cyanide plants, if so desired.

With the increasing severity and frequency of drought, the ability to modulate cyanide levels in preferred cassava cultivars will grow more important. TME 419, for example, is a popular cultivar in Nigeria known to be relatively low in cyanide (**Figure 4**), thus farming and preparation practices are likely tailored to these low levels. Under environmental conditions that would increase cyanogenesis, farmers and consumers using acyanogenic or reduced cyanide TME 419 would not have to alter these practices.

Reduction of cyanogen content in cassava has the potential for broad socioeconomic benefits for cassava producers and consumers as well as positive effects on the environment. As detoxification of cassava can take days (Tewe, 1992), acyanogenic cassava can reduce processing time and labor. Women and girls, who disproportionately bear the burden of this labor, may then be at greater liberty to pursue other forms of work and education. At the industrial scale, processing of acyanogenic cassava would not release cyanide into wastewater, thereby reducing the labor and cost of wastewater treatment and/or the toxicity to local terrestrial and aquatic life (Adewoye *et al*., 2005; Silva *et al*., 2017). Moreover, acyanogenic cassava cultivars would be a boon for food safety and consumer wellness. As described above, excessive consumption of cyanide with a protein-poor diet can lead to neurological harm including decline in motor proficiency and cognitive performance, and, in severe cases, paralysis (Kashala-Abotnes *et al*., 2019). Acyanogenic cassava could preclude these debilitating conditions and open at-risk consumers and their would-be caretakers to other pursuits.

Field testing of these edited lines and their corresponding wildtypes will allow well-controlled interrogation of the roles of cyanogens in cassava, including in herbivory defense and postharvest physiological deterioration. The editing approach we have taken can be applied to additional cassava varieties, notably, to those popular in regions of Africa projected to experience increased drought as a result of climate change and thus higher potential cyanide consumption. In addition to the *CYP79D* genes, the CRISPR-Cas9 system can be applied to modulate cyanogenic potential through modification of other genes of interest. A recent genome-wide association study, for example, identified an ATPase and MATE protein that regulate cassava cyanide levels in storage roots (Ogbonna *et al*., 2021). The potential for changing a particular trait in outbred cassava varieties, without disrupting the complement of other traits for which they are preferred, is an alluring aspect of this precision breeding method, and thus can play a role in the maintenance of genetic diversity across the global population of cassava cultivars.

## Experimental Procedures

### sgRNA and CRISPR construct design

Candidate target sequences were identified in *CYP79D1* and *CYP79D2* genes (Manes.13G094200 and Manes.12G133500, respectively, in cassava AM560-2 reference assembly v8.1, https://phytozome-next.jgi.doe.gov/info/Mesculenta_v8_1) of cassava using the online CRISPR-P 2.0 software (Liu *et al*., 2017). The software CasOT was used with default settings to search for potential off-targets in a 60444 genome assembly (Gomez *et al*., 2019; Xiao *et al*., 2014). Candidate targets with minimal off-target potential and approximately 500 bp apart were selected for assembly. Matching CRISPR targets in 60444 were verified by PCR amplification of targeted regions from genomic DNA extracts and Sanger sequencing.

The CRISPR-Cas9 expression entry plasmid (Thomazella *et al*., 2021) was re-engineered to carry the optimized gRNA scaffold with stem loop extension and A-U flip for improved Cas9 binding and gRNA transcription, respectively (Chen *et al*., 2014). The *BsaI* site in the backbone of the binary destination vector pCAMBIA2300 was removed via the QuikChange Site-Directed Mutagenesis Kit (Agilent) (Hajdukiewicz *et al*., 1994). The cassette carrying the CRISPR expression system was Gateway cloned into the *BsaI*-removed pCAMBIA2300 vector. The assembled binary vector with CRISPR expression system, pCAMBIA2300 CR3-EF, requires a single cloning step for insertion of desired gRNA with white colony screen and kanamycin selection in *E. coli*.

The selected CRISPR spacers were assembled into a polycistronic tRNA-gRNA (PTG) gene for multiplex targeting (Xie *et al*., 2015). The protocol was modified to incorporate the aforementioned stem loop extension and A-U flip. The Golden Gate cloning method was used to BsaI digest the pCAMBIA2300 CR3-EF vector and PTG ends, and then ligate the PTG into the vector. Sequences of assembled CRISPR constructs were verified via Sanger sequencing.

CRISPR construct activity was verified via *in planta* gemini-vector mutagenesis assay. Briefly, targeted regions were cloned into a derivative of the pLSL.D.R gemini-vector and pEAQ-HT vector maintaining replication elements for the generation of replicons bearing the target sites (Baltes *et al*., 2014; Sainsbury *et al*., 2009). Geminivirus constructs and CRISPR constructs were separately transformed into *Agrobacterium tumefaciens* strain GV3101 cultures via heat shock and rifampicin, gentamicin, and kanamycin selection. Transformants were grown overnight and diluted to OD600=0.3 each in infiltration medium (10 mM MES pH 5.6, 10 mM MgCl_2_, 150 µM acetosyringone). After incubation at room temperature for 3 h, *A. tumefaciens* cultures bearing CRISPR construct and corresponding geminivirus-targets construct were mixed 1:1 and infiltrated into *N. benthamiana* leaves. After five days, DNA was extracted from infiltrated leaf material via a modified CTAB procedure (Murray and Thompson, 1980). Frozen leaf tissue was ground by 3-mm glass beads in Minibeadbeater (Biospec Products, Inc.) and resuspended in extraction buffer (1.4 M NaCl, 100 mM Tris-HCl pH 8.0, 20 mM EDTA, 2% CTAB). Following incubation at 65 °C for at least 10 min, the extract was emulsified with chloroform and centrifuged at 16,000 *g* for 5 min. DNA was precipitated from the aqueous phase with an equal volume of isopropanol and centrifuged for 10 min at 4 °C. The supernatant was decanted and the DNA pellet was washed with 70% ethanol. After re-centrifugation for 2 min, ethanol was removed by pipette and air drying for 5–10 min. The DNA pellet was resuspended in 1X TE Buffer. Dissolution was advanced by incubation at 60–65 °C for 5 min, or overnight incubation at room temperature. Target regions were PCR amplified and run on 1.5% agarose gel. CRISPR-mediated excision of 500 bp between CRISPR targets resulted in an amplified band that was visibly smaller on the gel.

### Genetic transformation of cassava

*Agrobacterium*-mediated transformation was utilized to deliver CRISPR-Cas9 gene editing tools into friable embryogenic calli (FEC) of cassava accessions 60444 and TME 419, with subsequent plant regeneration, following the protocol described by Taylor *et al*. (Taylor *et al*., 2012) and Chauhan *et al*. (Chauhan *et al*., 2015). Somatic embryos were induced from leaf explants of *in vitro* micropropagated plants by culture on Murashige and Skoog basal medium (MS) (Murashige and Skoog, 1962) supplemented with 20 g/L sucrose (MS2) plus 50 µM picloram. Pre-cotyledon stage embryos were subcultured onto Gresshoff and Doy basal medium (GD) (Gresshoff and Doy, 1974) supplemented with 20 g/L sucrose and 50 µM picloram (GD2 50P) in order to induce production of FEC. Homogenous FEC were selected and used as target tissue for transformation with *Agrobacterium tumefaciens* strain LBA4404 (Taylor *et al*., 2012) or AGL1, carrying CRISPR constructs targeting *CYP79D1* (C1), *CYP79D2* (C2), and both genes (C3). In some cases infection was performed using sonication with a Branson 3510-DTH Ultrasonic Cleaner for three seconds. Transgenic tissues were selected and proliferated on GD2 50P containing paromomycin, prior to regeneration of embryos on MS2 medium supplemented with naphthalene acetic acid (NAA). Somatic embryos were germinated on MS2 medium containing 6-benzylaminopurine (BAP). Regenerated plants were maintained on MS2 in Phytatrays II (Sigma-Aldrich, St. Louis, MO), incubated at 28 °C in high light (90–150 μmol m^-2^ s^-1^ for 16 h light/8 h dark conditions) and subcultured every 3 weeks.

### Plant growth and maintenance

Plantlets were maintained in MS2 agar medium for stem elongation and stable growth. Well developed growing shoots were maintained in growth chambers in Phytatrays, one to two shoots per tray, following the conditions described by Taylor *et al*. (Taylor *et al*., 2012). Regenerated plants were micro-propagated and rooted in MS2 medium containing 2.2 g/L phytagel at two to three plantlets per petri dish. After three weeks rooted plantlets were transferred into soil (BM7 45% bark mix, Berger) in 3” square (0.37 L^3^) Kord pots and grown in a glasshouse. During soil transfer, pots were subirrigated with an aqueous solution containing, per gallon, Gnatrol WDG larvicide (Valent) at the label rate, ½ tsp Jack’s Professional LX 15-5-15 Ca-Mg fertilizer (J.R. Peters), ¼ tsp Jack’s Professional M.O.S.T. mix of soluble traces (J.R. Peters), and ½ tsp Sprint 330 (Becker Underwood). Plants were transferred to soil and watered, and the pots placed in trays with drainage holes. The trays were covered with a low (2”) dome and kept on a heating pad set to 80 °F in a misting bench for 100% humidity, under 40% white shade cloth. After 8–16 days, the low domes were replaced with 6” high vented domes and the trays moved into a room without shade, with misting 3 times per day. The domes were removed after 9–11 days and plants placed approximately six per tray in 28-pocket spacing trays. Some plants that were lagging were kept under non-vented high domes for longer periods. For the first four weeks after transfer to soil, plants were watered with Jack’s Peat Lite 15-16-17 (JR Peters) at 200 ppm approximately three times per week, and thereafter with Jack’s Blossom Booster 10-30-20 (JR Peters) at 100 ppm two times per week (Taylor *et al*., 2012). Plants were watered with tap water on days fertilizer was not administered.

### Sequence analysis

Putative transgenic lines (based on growth on antibiotic) were genotyped at the target loci. Genomic DNA extraction, PCR amplification, and Sanger sequencing were conducted as described in Gomez *et al*. (Gomez *et al*., 2019). PCR amplification and Sanger sequencing were performed using the gemini-CYP79D primers (**Table S2**).

Percent identity of the *CYP79D1* and *CYP79D2* promoter sequences and protein amino acid sequences was evaluated using sequences derived from the cassava v6.1 genome assembly (Bredeson *et al*., 2016). Sequences were aligned for analysis using Clustal Omega under default settings (Madeira *et al*., 2019). Protein domains were predicted using SMART (Letunic *et al*., 2021).

### Amplicon sequencing of selected CRISPR-Cas9 edited lines

For each line, leaf samples from two parts of an individual plant were collected for DNA extraction. PCR reactions for amplicon sequencing were performed using Phusion HF Polymerase and amplicon sequencing primers (**Table S2**), and amplified for 25 cycles. Samples were deep sequenced on an Illumina MiSeq using 300 bp paired-end reads to a depth of at least 10,000 reads per sample. Cortado (https://github.com/staciawyman/cortado) was used to analyze editing outcomes. Briefly, reads were adapter trimmed and then merged using overlap into single reads. These joined reads were then aligned to the target sequence using NEEDLE (Li *et al*., 2015) to identify any insertions or deletions (indels) overlapping the targeted cut site. Genotypes found in less than 1% of reads were considered to be PCR or sequencing errors.

### Off-target analysis

Identification of potential off-target loci was performed using CasOT software and a reference-based genome assembly of accession 60444, as described previously (Gomez *et al*., 2019; Xiao *et al*., 2014). Sites that contained the protospacer adjacent motif (PAM) region of the gRNA were selected and ranked according to sequence similarity to the target site, and the 2–3 highest ranking potential off-targets (those with the fewest mismatches to the guide) for each gDNA were selected for sequence analysis (**Table S1**). Genomic DNA was extracted from cassava leaves using the modified CTAB protocol described above. The selected potential off-target regions were amplified using Phusion polymerase (New England Biolabs [NEB]) and touchdown PCR (TD-PCR) (Korbie and Mattick, 2008), Phusion polymerase and 30 cycles of PCR with an annealing temperature of 63°C, or OneTaq Quick-Load 2X Master Mix with Standard Buffer (NEB) with 35 PCR cycles and an annealing temperature of 47 °C. PCR primer sequences are listed in **Table S2**. The TD-PCR protocol began with an annealing temperature of Tm + 10 °C for the first cycling phase (Tm calculated by NEB Tm Calculator tool). The annealing temperature was then decreased by 1 °C per cycle until the primers’ Tm was reached, followed by 20 or 25 cycles using the primer Tm as the annealing temperature. PCR amplicons were visualized using gel electrophoresis, then the remaining reaction was purified for sequencing via the AccuPrep PCR/Gel Purification Kit (Bioneer), the Monarch PCR & DNA Cleanup Kit (NEB), or SPRI magnetic nucleic acid purification beads (UC Berkeley DNA Sequencing Facility). We sequenced purified amplicons containing potential off-targets using Sanger sequencing. Putative off-target loci were then examined in SnapGene for potential sequence discrepancy with the 60444 reference sequence.

### RT-PCR

Cassava cDNA was generated using the Spectrum Plant Total RNA Kit (Sigma-Aldrich) following Protocol A and performing On-Column DNase Digestion. Concentration of RNA extracts were measured by NanoDrop One (Thermo Fisher Scientific). Quality of RNA was examined by first denaturing aliquots at 70 °C for 5 min (followed by 4 °C on ice), then electrophoresing 200 ng of RNA on 1.5% UltraPure Agarose (Invitrogen). 450–1000 ng of RNA was added to SuperScript III Reverse Transcriptase (Invitrogen) reaction mix with Oligo(dT)_20_. Reaction was run for 60 minutes at 50 °C followed by RNase H treatment. Primers were designed to amplify the *CYP79D* transcripts from the 5’ UTR to the 3’ UTR. 2 µL of cDNA mix was added to 50 µL Phusion High-Fidelity DNA Polymerase (NEB) reaction mix. cDNA was amplified for 35 cycles. PCR reactions were run on 1.5% agarose and desired bands were extracted. Amplicons were cloned into the Zero Blunt PCR Cloning Kit (Thermo Fisher Scientific) and 10–12 colonies subsequently sequenced via the UC Berkeley DNA Sequencing Facility.

### Measurement of cyanogenic potential

#### Measurement of cyanogens from *in vitro* plantlets

We used LC-MS to measure linamarin and lotaustralin in *in vitro* plantlets. Regenerated transgenic plants were micro-propagated and established in MS2 medium at two plantlets per Phytatray in a growth chamber at 28 °C +/- 1 °C, 41% relative humidity, 120–150 μmol/m^2^/s light for 16 h light/8 h dark conditions. After four to five weeks plants were ready for tissue sampling. One leaf was harvested from each plantlet and stored in a plastic bag on ice until extraction, approximately 1–3 h later. For biological replicates, we harvested tissue from three plantlets per line.

Approximately 20 or 30 mg of leaf tissue was excised from fresh leaves and placed in a 1.5-mL tube (Safe-Lock, Eppendorf) with 600 or 900 µL of 85% MeOH warmed to approximately 68 °C. Sample weight was recorded. Negative controls contained no tissue. A cap lock was added and the tube was floated in boiling water for three minutes, then returned to ice. One to three tubes were boiled at a time. Cooled tubes were spun down briefly. A 1:10 or 1:20 dilution was prepared from each extract, pipetted up and down to mix, and spun through a 0.45-µM spin filter (Ultrafree MC HV Durapore PVDF, EMD Millipore) for 2 min, 10,000 x *g*, 4 °C. 20 µL of filtered extract was placed in a glass autosampler vial with insert (Fisher Scientific), and the LC-MS run begun the same day. Three samples were submitted from each extract, for technical replicates.

To facilitate absolute quantitation, standard stocks were prepared from solid linamarin (Cayman Chemical, purity ≥98%) and lotaustralin (Millipore Sigma, purity ≥95%) resuspended in 85% MeOH to 3 or 4 mM, aliquoted into dark glass vials, and stored at –20 °C. On the day of assay, these standards were further diluted in 85% MeOH and submitted for LC-MS analysis. Lotaustralin standards ranged in concentration from 0.01 to 1 µM, and linamarin from 0.05 to 5 µM. Submitted standard samples contained both linamarin and lotaustralin in known quantities. As with extracts, three technical replicates were performed for each standard. To buffer against any potential position/timing effects, samples were analyzed by LC-MS in three consecutive cohorts, where each cohort had one technical replicate from each sample.

Samples of cassava extracts were analyzed using a liquid chromatography (LC) system (1200 series, Agilent Technologies, Santa Clara, CA) that was connected in line with an LTQ-Orbitrap-XL mass spectrometer equipped with an electrospray ionization (ESI) source (Thermo Fisher Scientific, San Jose, CA). The LC system contained the following modules: G1322A solvent degasser, G1311A quaternary pump, G1316A thermostatted column compartment, and G1329A autosampler (Agilent). The LC column compartment was equipped with a reversed-phase analytical column (length: 150 mm, inner diameter: 1.0 mm, particle size: 5 µm, Viva C18, Restek, Bellefonte, PA). Acetonitrile, formic acid (Optima LC-MS grade, 99.5+%, Fisher, Pittsburgh, PA), and water purified to a resistivity of 18.2 MΩ·cm (at 25 °C) using a Milli-Q Gradient ultrapure water purification system (Millipore, Billerica, MA) were used to prepare LC mobile phase solvents. Solvent A was 99.9% water/0.1% formic acid and solvent B was 99.9% acetonitrile/0.1% formic acid (volume/volume). The elution program consisted of isocratic flow at 2% B for 2 min, a linear gradient to 6% B over 1 min, a linear gradient to 90% B over 0.5 min, isocratic flow at 90% B for 4.5 min, a linear gradient to 2% B over 0.5 min, and isocratic flow at 2% B for 16.5 min, at a flow rate of 200 µL/min. The column compartment was maintained at 40 °C and the sample injection volume was 10 µL. Full-scan mass spectra were acquired in the positive ion mode over the range of mass-to-charge ratio (*m*/*z*) = 200 to 800 using the Orbitrap mass analyzer, in profile format, with a mass resolution setting of 100,000 (at *m*/*z* = 400, measured at full width at half-maximum peak height, FWHM). For tandem mass spectrometry (MS/MS or MS^2^) analysis, selected precursor ions were fragmented using collision-induced dissociation (CID) under the following conditions: MS/MS spectra acquired using the linear ion trap, in centroid format, normalized collision energy: 35%, activation time: 30 ms, and activation Q: 0.25. Mass spectrometry data acquisition and analysis were performed using Xcalibur software (version 2.0.7, Thermo Fisher Scientific).

To convert LC-MS concentration values in µM to grams per kg fresh weight (fw), the following formula was used:

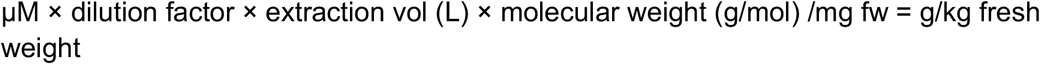

where the molecular weight of linamarin is 247.248 g/mol and of lotaustralin is 261.272 g/mol. Concentration values reported as <LLOQ (below the lower limit of quantification) were treated as 0 µM.

#### Measurement of cyanide from cassava plants growing in glasshouse

Each assay cohort was grown from *in vitro* plantlets transferred to soil on the same day. Plants were collected from the greenhouse six to eight months after transfer to soil. The height of each plant’s stem(s) were measured from the topsoil to the apical meristem in a direct line, without forcing the stems to bend. Leaf and tuberous root samples were collected for cyanide content analysis via the picrate paper assay using Konzo Kit A from Australia National University Konzo Prevention Group ((Bradbury *et al*., 1999); https://biology.anu.edu.au/research/resources-tools/konzo-kits). Leaf samples were collected from three plants of each line. The third, fourth, and fifth expanded leaves from the top were cut perpendicular to the midribs into 0.5-cm wide pieces with clean scissors. Leaf cuts were immediately ground with a mortar and pestle. One hundred milligrams of ground leaf was loaded into a vial containing buffer paper, and 1 mL of water and the cyanide indicator paper were added immediately and the vial capped. For negative controls, no tissue was added to a vial. Positive controls were conducted using the standard provided with the kit. Sample vials were incubated overnight (minimum 12 h) at room temperature. Up to nine tuberous roots were collected from each line, with no more than three roots coming from a single plant. To buffer against any cyanide variation over the course of processing samples, root collection was staggered among groups to three to four roots per line at a time. Roots of minimum 1 cm in diameter were collected, washed, and photographed by a ruler for scale. Each root was cut at its widest section, and a 1.5-mm slice was taken crosswise via a kitchen mandoline. The peel (rind) was removed and 100 mg of tuberous root loaded into a vial and sealed as described above.

Indicator papers were removed from vials and compared to a cyanide color chart for an approximate cyanide content reference. These papers were then placed in 15 mL culture tubes and completely immersed in 5 mL of water. Solutions were incubated at room temperature for 30–60 minutes with occasional gentle stirring. The absorbance of each pipette-mixed solution was measured at 510 nm using an Ultrospec 3000 UV/Visible Spectrophotometer (Pharmacia Biotech). Absorbance was normalized to the value of the negative control (no plant sample). Absorbance was multiplied by 396 to acquire the total cyanide content in ppm (equivalent to mg HCN per kg of tissue).

#### Statistical analyses

Box and whisker plots were generated for cyanide measurements from picrate assays. The box of each plot represents the interquartile range (IQR) which is bounded by a lower quartile (Q1, 25th percentile) and an upper quartile (Q3, 75th percentile) of the data. The whiskers of each plot are defined as the approximate minima and maxima of the data, with the minimum data value defined as Q1 – 1.5*IQR, and the maximum defined as Q3 + 1.5*IQR. Data outside of the maximum and minimum values were considered outliers.

The Wilcoxon rank-sum statistic (also known as the Mann–Whitney *U* test) was performed on the picrate data to test whether there were statistically significant differences between the various lines, using pairwise comparisons. This method was used primarily due to the nonparametric and continuous nature of our picrate data. Two-group Wilcoxon rank-sum statistical comparisons between all lines were performed via SciPy’s stats.ranksum function (https://docs.scipy.org/doc/scipy/reference/generated/scipy.stats.ranksums.html), which uses a normal approximation of the rank sum. The p-values of each two-group test were compiled to form heatmaps. In these heatmaps, p-values less than 0.05 were considered significant and shaded blue, indicating that for two compared groups, the data appeared to come from two separate distributions. P-values greater than 0.05 were shaded red and indicated that for two compared groups, the data appeared as if drawn from the same distribution, demonstrating statistical insignificance from each other.

All plots were generated in Google Colab notebooks with the Seaborn data visualization library.

## Supporting information

File S1: Genotypes of CRISPR/Cas9-edited 60444 and TME 419 lines

File S2: Predicted CYP79D amino acid sequences from selected 60444 and TME 419 mutant lines

File S3: LC-MS and picrate readings

File S4: Python notebook for figs 2-4

## Author Contributions

Designed the project: MAG, NJT, BJS, M-JC, DSR, JBL.

Molecular biology lead: MAG.

Gemini-vector transient assay: MAG, AS.

Cassava transformation: BG, RDC.

Genotyping of on- and off-target sites: MAG, KCB, SEL, JMM.

Extractions for LC-MS: MAG, BG, JBL.

LC-MS: ATI.

LC-MS plots: JBL.

Picrate assays: MAG, KCB, SEL, JBL.

Picrate assay plots and statistical analysis: KCB.

Illumina amplicon data analysis: SW.

Project leadership: JBL, with co-PIs BJS, M-JC, DSR.

Wrote the paper: MAG and JBL; with contributions by KCB, BG, ATI, SEL, SW, NJT, M-JC; and edited by NJT, M-JC, and DSR.

All authors reviewed and approved the manuscript.

## Acknowledgements

The IGI is grateful to the Danforth Center for support in initiating its cassava transformation and regeneration platform, including Tira Jones and Danielle Stretch for technical assistance, and Claire Albin for sharing plant care protocols. The authors thank Tina Wistrom and UCB Oxford Tract greenhouse staff for plant care; Jonathan Vu and Netravathi Krishnappa, IGI Center for Translational Genomics, for Illumina Amplicon sequencing; UC Berkeley DNA Sequencing Facility; Elaine Zhang, Dominick Tucker, Xiuli Shen, and Ankita Singh for technical assistance; and Benton Cheung for cassava graphic. We are grateful to Ros Gleadow for advice on cyanide measurements, Kirsten Jørgensen for sharing the extraction protocol for LC-MS, John Young and Nikki Kong for advice on statistical analyses, and Susan Abrahamson for consultation on intellectual property.

This work was supported by the Innovative Genomics Institute. The QB3/Chemistry Mass Spectrometry Facility received support from the National Institutes of Health (grant 1S10OD020062-01).

## Availability of data & materials

Vectors are available upon request to BJS. Plants are available upon request to JBL.

## Supporting Information

**Figure S1** CRISPR-Cas9 construct activity assay via targeting of surrogate gemini-vector.

**Figure S2** Cas9 expression system.

**Figure S3** CRISPR-Cas9 induces indels at *CYP79D1* and *CYP79D2* gRNA target sites in transgenic TME 419 lines.

**Table S1** Putative off-target loci in 60444.

**Table S2** PCR primers used in this work.

**Table S3** Off-target results for 60444.

**Figure S4** Representative 6044 edited plants and storage roots

**Figure S5** Cyanogen levels in edited 60444 *in vitro* plantlets.

**Figure S6** Cyanogen levels in edited TME 419 *in vitro* plantlets.

**Figure S7** Sanger and Illumina sequence analysis of TME 419 line C3-10 at target site 2B.

**Figure S8** Tissue-specific transcript expression of *CYP79D1* and *CYP79D2*.

**File S1** Genotypes of CRISPR/Cas9-edited 60444 and TME 419 lines.

**File S2** Predicted CYP79D amino acid sequences from selected 60444 and TME 419 mutant lines.

**File S3** LC-MS and picrate readings.

**File S4** Python notebook for figures 2–4.

