## Supporting Information for "CRISPR-Cas9-mediated knockout of *CYP79D1* and *CYP79D2* in cassava attenuates toxic cyanogen production"

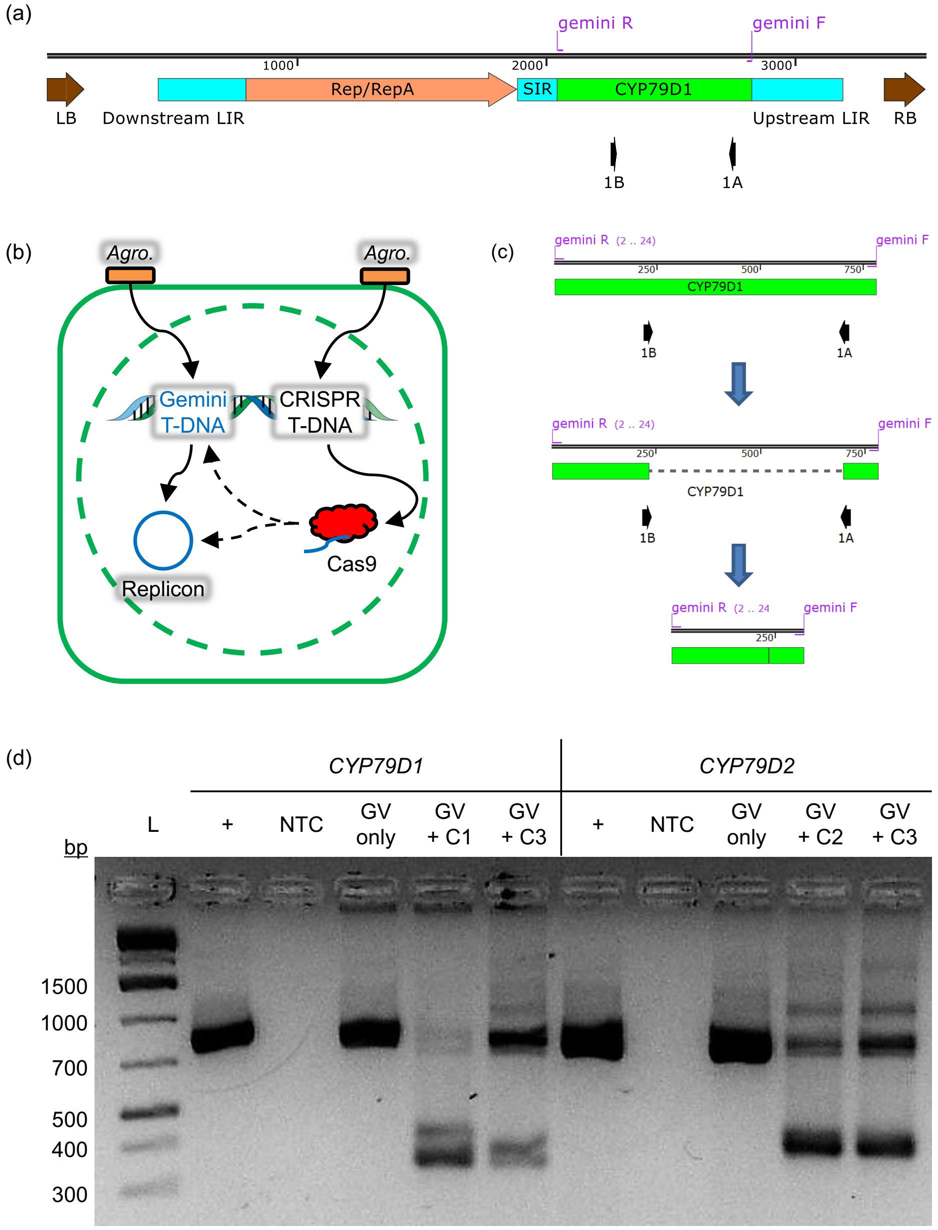


**Figure S1** CRISPR-Cas9 construct activity assay via targeting of surrogate gemini-vector. (a) A 781-bp fragment of *CYP79D1* was cloned within the replication sequences essential for geminiviral replicon synthesis. CRISPR-Cas9 target sites 1A and 1B are within this *CYP79D1* fragment. The replication initiator protein (Rep) initiates rolling-circle replication by binding to the large intergenic region (LIR). Replicons will include the *CYP79D1* fragment. Primer sequences for the amplification of the *CYP79D1* fragment are shown as “gemini F" and “gemini R.” (b) The assembled gemini-vector and CRISPR-Cas9 vector encoding the respective gRNA are co-agroinfiltrated into *Nicotiana benthamiana* leaf tissue (nuclear membrane shown as dashed green line). Following expression of the CRISPR-Cas9 system and replicon synthesis, Cas9-gRNA complexes may target either the gemini-vector T-DNA or replicons themselves (paths shown as dashed black lines). Mutagenesis of CRISPR-Cas9 target sites is proliferated in subsequently generated replicons. (c) Simultaneous CRISPR-Cas9 mediated cutting of the target sites can cause excision of the intervening sequence. Amplification by primers “gemini F” and “gemini R” now yields a 361-bp fragment. The same approach shown in (a–c) was taken with *CYP79D2*, using a 784-bp fragment encompassing target sites 2A and 2B. (d) DNA is extracted from leaves infiltrated with the gemini-vector and with/without the respective CRISPR-Cas9 construct (C1, C2, C3). Gel electrophoresis of PCR amplicons shows band sizes of wholly intact and excised DNA fragments. Wholly intact DNA fragments are expected to be 781 bp whereas excised DNA fragments are expected to be 361 bp. +, gemini-vector carrying intact *CYP79D* fragment; NTC, no template control; GV, gemini-vector. Maps created with SnapGene.

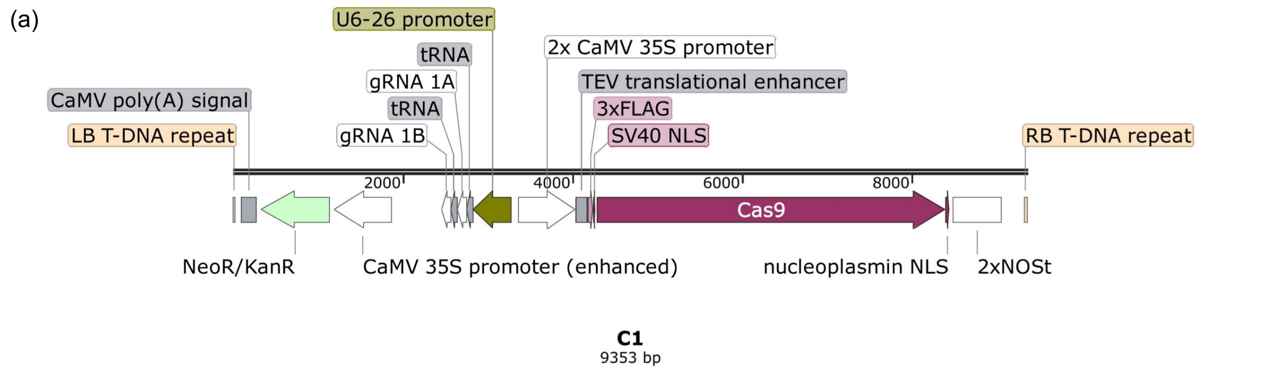


**Figure S2** Cas9 expression system. 2x35S promoter drives expression of human codon optimized Cas9, preceded by TEV translational enhancer. *Arabidopsis thaliana* U6-26 promoter drives expression of polycistronic tRNA-gRNAs. (a) C1 T-DNA, coding for two gRNAs targeting *CYP79D1*. (b) C2 T-DNA, coding for two gRNAs targeting *CYP79D2.* (c) C3 T-DNA, including four total gRNAs, two each targeting *CYP79D1* and *CYP79D2*. LB, left border; NeoR/KanR, Neomycin/Kanamycin resistance; T-DNA, transfer DNA; gRNA, guide RNA; tRNA, transfer RNA; CaMV, Cauliflower Mosaic Virus; TEV, Tobacco Etch Virus; SV40, Simian Virus 40; NLS, nuclear localization sequence; NOSt, nopaline synthase terminator; RB, right border. Maps created with SnapGene.

(a)

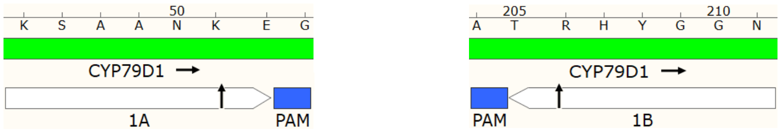


| Line | 1A sequence | Frame shift | 1B sequence | Frame shift |
|---|---|---|---|---|
| WT | AAGAGTGCTGCTAACAA GGAAGG | × | CCACCA GGCATTACGGCGGGAAT | × |
| C1-2 | AAGAGTGCTGCTAACAATGGAAGG | ✓ | CCACCA GGCATTACGGCGGGAAT<br>CCACCA -GCATTACGGCGGGAAT | ×<br>✓ |
| C1-9 | AAGAGTGCTGCTAACAAAGGAAGG | ✓ | CCA--- ----TTACGGCGGGAAT<br>CCACCAAGGCATTACGGCGGGAAT | ✓<br>✓ |
| C1-10 | AAGAGTGCTGCTAACAAAGGAAGG<br>AAGAGTGCTGCTAA--A GGAAGG | ✓<br>✓ | CCACCAGGGCATTACGGCGGGAAT<br>CCACCA -GCATTACGGCGGGAAT | ✓<br>✓ |
| C3-4 | AAGAGTGCTGCTAAC-- ------...<br>AAGAGTGCTGCTAAC-- --AAGG | <br>✓ | ...------ -GCATTACGGCGGGAAT<br>CCACCA GGCATTACGGCGGGAAT | ✓<br>× |

(b)

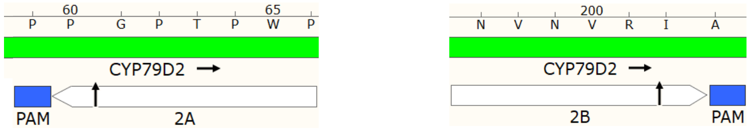


| Line | 2A sequence | Frame shift | 2B sequence | Frame shift |
|---|---|---|---|---|
| WT | CCCTCCT GGTCCGACTCCATGGCC | × | GAATGTGAATGTGAGAA TTGCGG | × |
| C2-6 | CCCTCCTAGGTCCGACTCCATGGCC<br>CCC---- -GTCCGACTCCATGGCC | ✓<br>✓ | GAATGTGAA-------- ------*<br>GAATGTGAATGTGAGAAATTGCGG | ✓<br>✓ |
| C2-7 | CCCTCCT ----CGACTCCATGGCC<br>CCCTCCT -------CTCCATGGCC | ✓<br>✓ | GAATGTGAATGTG---- TTGCGG<br>GAATGTGAATGTGAGAATTTGCGG | ✓<br>✓ |
| C2-10 | CCCTCCTAGGTCCGACTCCATGGCC<br>CCCTCCT -GTCCGACTCCATGGCC | ✓<br>✓ | GAATGTGAATGTGAGA- ----GG<br>GAATGTGAATGTGA--A TTGCGG | ✓<br>✓ |
| C3-4 | C------ --------------GCC<br>CCCTCCT GGTCCGACTCCATGGCC | ✓<br>× | GAATGTGAATGTGAGAAATTGCGG<br>GAATGTGAATGTGAGAATTTGCGG | ✓<br>✓ |

**Figure S3** CRISPR-Cas9 induces indels at *CYP79D1* and *CYP79D2* gRNA target sites in transgenic TME 419 lines. Diagrams of the protospacers (white) and protospacer adjacent motifs (PAMs, blue) of *CYP79D1* (a) and *CYP79D2* (b) gRNA targets are aligned to edited line genotypes. Edited lines are identified by the CRISPR construct with which they were modified (C1, C2, C3), followed by an index number. Black arrow indicates predicted CRISPR-Cas9 cut site. Lengths are to amino acid (top bar) and nucleotide (bottom table) scale. Homozygous genotypes are shown as a single sequence per line. Bi-allelic genotypes are shown as two sequences per line. Multiple mutations to a single allele are row-matched. Insertions are denoted by red, underlined nucleotides. Deletions are denoted by red dashes. Presence of a frameshift mutation at the corresponding target site is denoted by ✓; absence of a frameshift mutation is denoted by ×. Asterisk denotes a deletion larger than shown: line C2-6 has a 37 bp deletion. Maps created with SnapGene.

**Table S1** Putative off-target loci in 60444**.** Highest ranking off-targets identified by CasOT utilizing the 60444 genome assembly (Gomez *et al.*, 2019). Lower case letters in the Sequence column identify mismatches with the target sequence. Non-seed and seed regions separated by underscore. Spacer and PAM separated by hyphen. Locations and gene identifiers provided are for cassava AM560-2 v6.1 reference genome.

| Gene | Target | # | Sequence | Number of mismatches (Non-seed, seed) | V6.1 Location | Region | Gene Identifier |
|---|---|---|---|---|---|---|---|
| CYP79D1 | 1A | 1 | `tAaAaaca_TGCTAACAAGGA-TGG` | 6, 0 | Chromosome04: 3401297..3401319 | intergenic region | - |
| | | 2 | `AgagtaGt_TGCTAACAAGGA-TGG` | 6, 0 | Chromosome08: 1654375..1654397 | intron | Manes.08G 017500 |
| | 1B | 1 | `AgTggCcC_CGTgATGCCTGG-TGG` | 4, 1 | Chromosome05: 4248805..4248827 | intergenic region | - |
| | | 2 | `ATatCgGa_CGTgATGCCTGG-CGG` | 4, 1 | Chromosome08: 4989884..4989906 | intergenic region | - |
| CYP79D2 | 2A | 1 | `GaCttaGG_AGTCtGACCAGG-AGG` | 4, 1 | Chromosome02: 18109801..18109827 | intergenic region | - |
| | | 2 | `GGggAatG_AGTCGGACgAGG-TGG` | 4, 1 | Chromosome04: 17588060..17588082 | exon | Manes.04G 064300 |
| | | 3 | `ttCttTGG_AGTCGGAgCAGG-AGG` | 4, 1 | Chromosome15: 10501754..10501776 | intergenic region | - |
| | 2B | 1 | `cccTaaGA_ATGTGAGAATTG-AGG` | 5, 0 | Chromosome03: 17132076..17132098 | intron | Manes.03G 098600 |
| | | 2 | `GgtTtgtA_ATGTGAGAATTG-AGG` | 5, 0 | Chromosome15: 9415632..9415654 | intergenic region | - |

**Table S2** PCR primers used in this work.

| Name | Sequence | Function |
|---|---|---|
| CYP79D-1 TR F | CAACACGGTCAAGATCTTGTTCG | preliminary seq analysis of target region |
| CYP79D-2 TR F | GAACAATACTGCCAAAATCCTCC | preliminary seq analysis of target region |
| CYP79D TR R | ATCCCTTATGGTCTTATTTGCAT | preliminary seq analysis of target region |
| gemini-CYP79D1 TR F | AATTATTCGTACGACCCTCCCAACACGGTCAAGATCTTG | seq analysis and gemini-vector cloning |
| gemini-CYP79D2 TR F | AATTATTCGTACGACCCTCCGAACAATACTGCCAAAATC | seq analysis and gemini-vector cloning |
| CYP79D-gemini TR R | CATAAAATAATCATTTTATTCATCCCTTATGGTCTTATTTG | seq analysis and gemini-vector cloning |
| CYP79D1 g1A-F | TAGGTCTCCTGCTAACAAGGAGTTTAAGAGCTATGC | gRNA assembly into PTG array |
| CYP79D1 g1A-R | ATGGTCTCAAGCAGCACTCTTTGCACCAGCCGGGAA | gRNA assembly into PTG array |
| CYP79D1 g1B-F | TAGGTCTCCCGTAATGCCTGGGTTTAAGAGCTATGC | gRNA assembly into PTG array |
| CYP79D1 g1B-R | ATGGTCTCATACGGCGGGAATTGCACCAGCCGGGAA | gRNA assembly into PTG array |
| CYP79D2 g2A-F | TAGGTCTCCAGTCGGACCAGGGTTTAAGAGCTATGC | gRNA assembly into PTG array |
| CYP79D2 g2A-R | ATGGTCTCAGACTCCATGGCCTGCACCAGCCGGGAA | gRNA assembly into PTG array |
| CYP79D2 g2B-F | TAGGTCTCCATGTGAGAATTGGTTTAAGAGCTATGC | gRNA assembly into PTG array |
| CYP79D2 g2B-R | ATGGTCTCAACATTCACATTCTGCACCAGCCGGGAA | gRNA assembly into PTG array |
| Ampseq-CYP79D1-g1A F | GCTCTTCCGATCTCCACCATCGGTTTACTTAACG | MiSeq analysis |
| Ampseq-CYP79D1-g1A R | GCTCTTCCGATCTAACAAAGTTAGTTCTTCCAAAACG | MiSeq analysis |
| Ampseq-CYP79D1-g1B F | GCTCTTCCGATCTTCTTAACCTCAGAGATCATTTCTCC | MiSeq analysis |
| Ampseq-CYP79D1-g1B R | GCTCTTCCGATCTGAAAGGCAAGAAATCTGATATGC | MiSeq analysis |
| Ampseq-CYP79D2-g2A F | GCTCTTCCGATCTCGTCCTCCATGAACAATACTGC | MiSeq analysis |
| Ampseq-CYP79D2-g2A R | GCTCTTCCGATCTATTTCACGAGCAATGACAGG | MiSeq analysis |
| Ampseq-CYP79D2-g2B F | GCTCTTCCGATCTTCCTAACTTCAGAGATCATTTCTCC | MiSeq analysis |

| | | |
|---|---|---|
| Ampseq-CYP79D2-g2B R | GCTCTTCCGATCTAAAAGGCAAGTAATCAGAGATGC | MiSeq analysis |
| iSeq - CYP79D1 g1A F | GCTCTTCCGATCTCACCTCCTTCGCCTCCTC | iSeq analysis |
| iSeq - CYP79D1 g1A R | GCTCTTCCGATCTTGAGTTGGTGAATCCACCG | iSeq analysis |
| iSeq - CYP79D1 g1B F | GCTCTTCCGATCTGCTAGACACAAATGGCTCCATG | iSeq analysis |
| iSeq - CYP79D1 g1B R | GCTCTTCCGATCTACGGCATCAATGTGCTCG | iSeq analysis |
| iSeq - CYP79D1 g1C F | GCTCTTCCGATCTTTGGCGATATCCCTGGATTTG | iSeq analysis |
| iSeq - CYP79D1 g1C R | GCTCTTCCGATCTGGAGGGAGTGGGAGTTTC | iSeq analysis |
| iSeq - CYP79D2 g2A F | GCTCTTCCGATCTTACTGCCAAAATCCTCCTTATCAC | iSeq analysis |
| iSeq - CYP79D2 g2A R | GCTCTTCCGATCTTCAGACAAATATCGGTGTTCATGTC | iSeq analysis |
| iSeq - CYP79D2 g2B F | GCTCTTCCGATCTATTTCTCCAGCTAGGCACAAATG | iSeq analysis |
| iSeq - CYP79D2 g2B R | GCTCTTCCGATCTACGTGCATGATTTCTTCAGG | iSeq analysis |
| iSeq - CYP79D2 g2C F | GCTCTTCCGATCTGGACTTGAATTGTTTAGGGCAAC | iSeq analysis |
| iSeq - CYP79D2 g2C R | GCTCTTCCGATCTCCATGGAGTCGGACCAGG | iSeq analysis |
| 1A OT1 F | GCAAGTTGCATGGAAGTCTCTC | off target analysis |
| 1A OT1 R | TCCTTCTTACATGCTTCTCAAGG | off target analysis |
| 1A OT2 F | AGGACAGAAATGGAATGATGC | off target analysis |
| 1A OT2 R | TATCCAAGAGGGGCTCGTAG | off target analysis |
| 1B OT1 F | TGCAAAACATGTCAGCTCAAC | off target analysis |
| 1B OT1 R | TCACCACATATACTGCCTTTGC | off target analysis |
| 1B OT2 F | GCCGGGTATTACATCCTTCC | off target analysis |
| 1B OT2 R | ATGCCAAGTCACAAGGTGAG | off target analysis |
| 2A OT1 F | ATGCCTCTCCGCTATAGGAC | off target analysis |
| 2A OT1 R | AGTTTGATCATGCTAAATGAAGG | off target analysis |

| | | |
|---|---|---|
| 2A OT2 F | TTGATATCTATTATTCGTTTTCTGGAC | off target analysis |
| 2A OT2 R | TCATCAATTGCAAGGCTCTTC | off target analysis |
| 2A OT3 F | AAATAGTCATTTCGTCTATTTTTGC | off target analysis |
| 2A OT3 R | ACGAAATAGTCCTTCCTCATCG | off target analysis |
| 2B OT1 F | GAATTAGGGAGGAAATGACAAAAG | off target analysis |
| 2B OT1 R | CACCATTTCTTCTTGCAAAGC | off target analysis |
| 2B OT2 F | ATTTTCTTTTTCAATTCTAACTTCAAC | off target analysis |
| 2B OT2 R | GGCACATGCGACTTCTGTG | off target analysis |
| CYP79D1 5UTR F | GCGATATCCCTGGATTTG | cDNA analysis |
| CYP79D1 3UTR R | ATTAAAGGACGTTCTAAGAAC | cDNA analysis |
| CYP79D2 5UTR F | GTATGGTCTTGGTCATAGC | cDNA analysis |
| CYP79D2 3UTR R | CTAACAACTCACATTCATCC | cDNA analysis |

**Table S3** Off-target results for 60444. Highest ranked potential off-targets (O.T.) were examined by Sanger sequence analysis for conservation of the wildtype (WT) sequence. Check marks indicate sequences from the WT line matched the 60444 assembly. Edited lines are identified by the CRISPR construct with which they were modified (C1, C2, C3), followed by an index number (e.g. 142A). Corresponding loci in edited lines that matched the WT sequence are marked as WT. Loci that could not be amplified and sequenced are marked as not determined (N.D.).

| **Gene** | **Target** | **O.T.** | **WT** | **C1-4** | **C1-6** | **C2-4** | **C2-2** | **C3-2** | **C3-142A** | **C3-145C** |
|---|---|---|---|---|---|---|---|---|---|---|
| *CYP79D1* | 1A | 1 | ✓ | WT | WT | | | WT | WT | WT |
| | | 2 | ✓ | WT | WT | | | WT | WT | WT |
| | 1B | 1 | ✓ | WT | WT | | | WT | WT | WT |
| | | 2 | ✓ | WT | WT | | | WT | WT | WT |
| *CYP79D2* | 2A | 1 | ✓ | | | N.D. | N.D. | N.D. | N.D. | N.D. |
| | | 2 | ✓ | | | WT | WT | WT | WT | WT |
| | | 3 | ✓ | | | WT | WT | WT | WT | WT |
| | 2B | 1 | ✓ | | | WT | WT | WT | WT | WT |
| | | 2 | ✓ | | | WT | WT | WT | WT | WT |

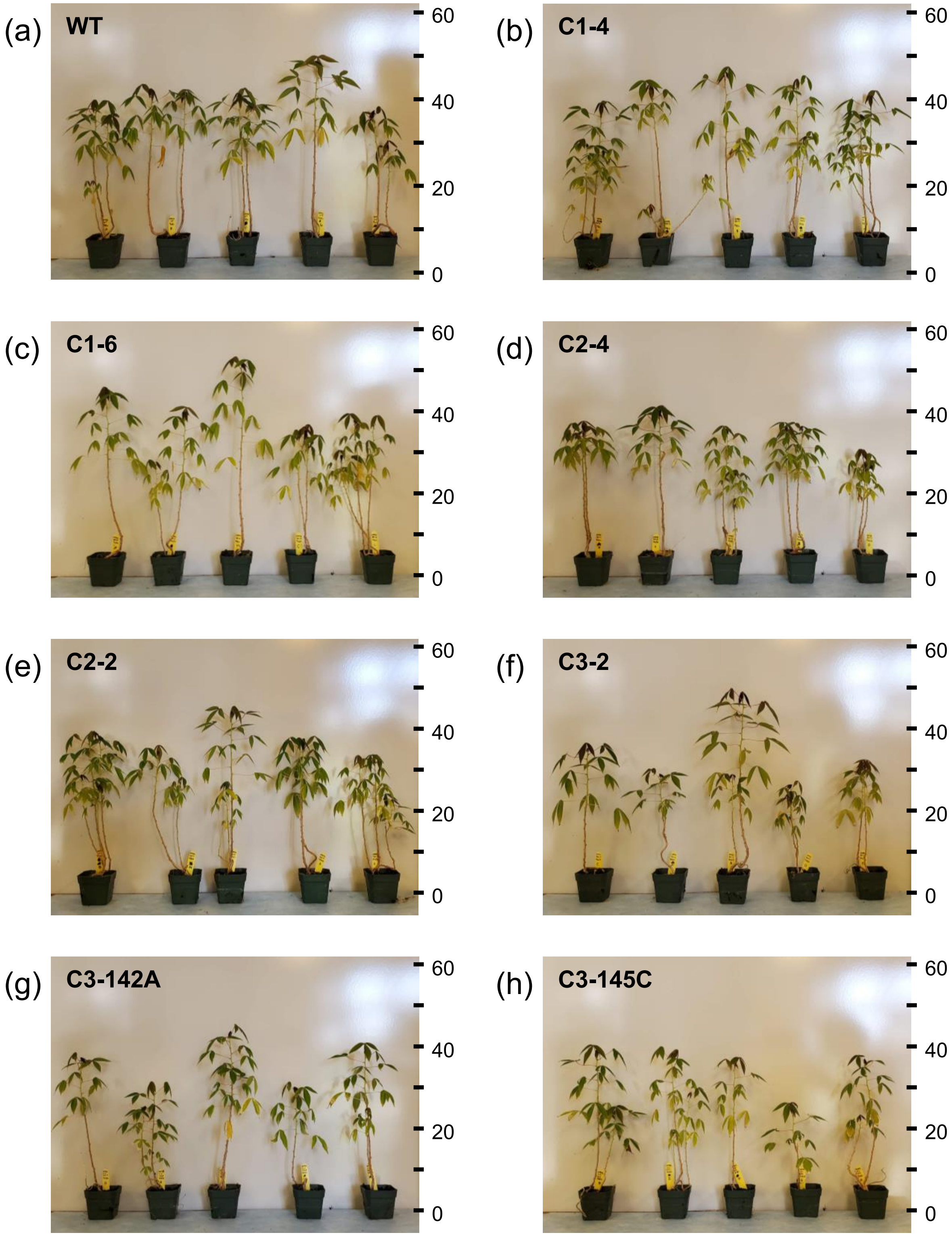
(a)
WT
(b)
C1-4
(c)
C1-6
(d)
C2-4
(e)
C2-2
(f)
C3-2
(g)
C3-142A
(h)
C3-145C
60
40
20
0

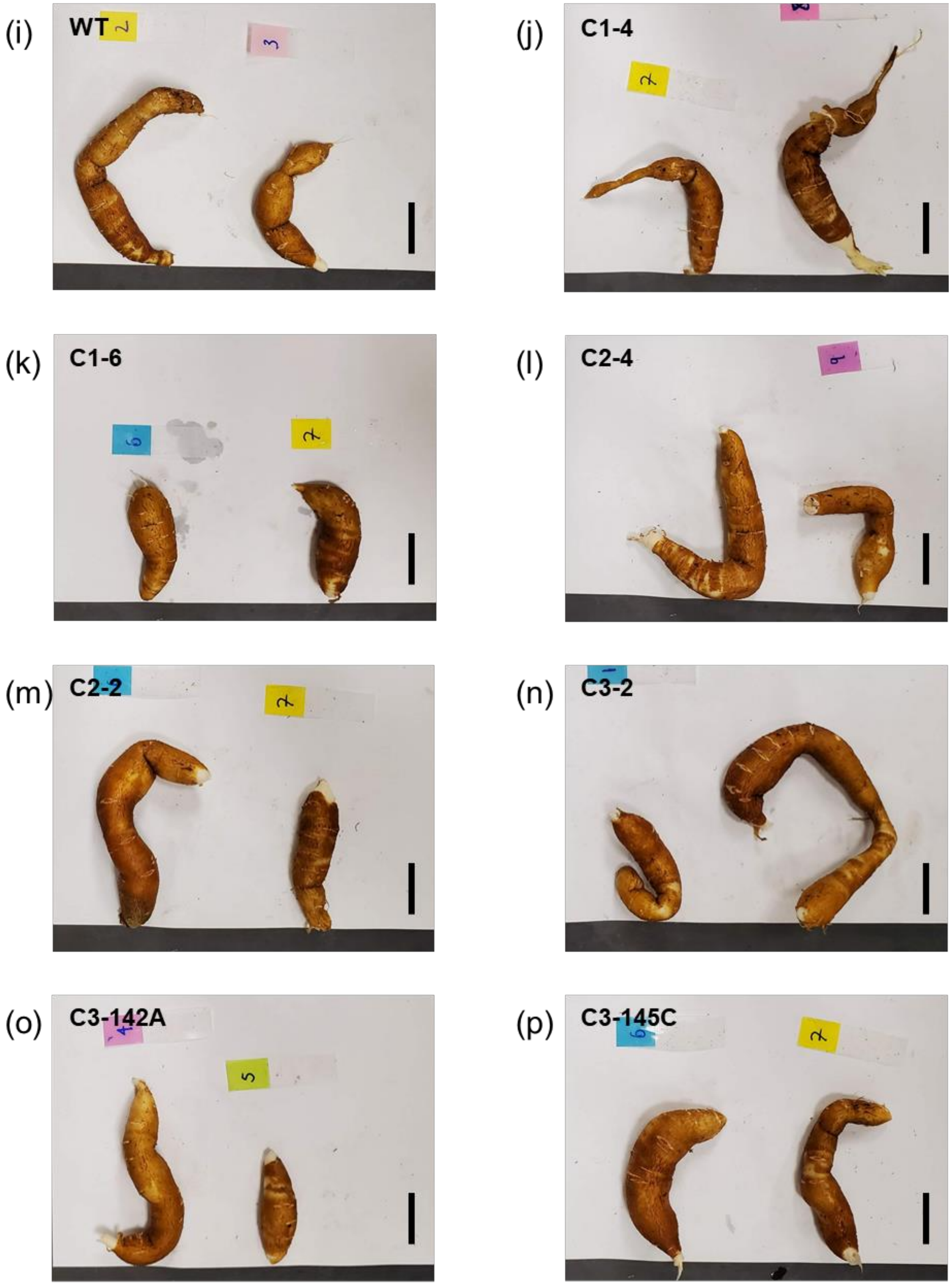


**Figure S4** Representative 60444 genome edited plants and storage roots. (a–h) Plants growing in 3-inch pots eight months after transfer to soil. Scale is in centimeters. (i-p) Roots harvested from plants grown in 3-inch pots 6.5 months after transfer to soil. Black bar represents 2 cm.

(a)

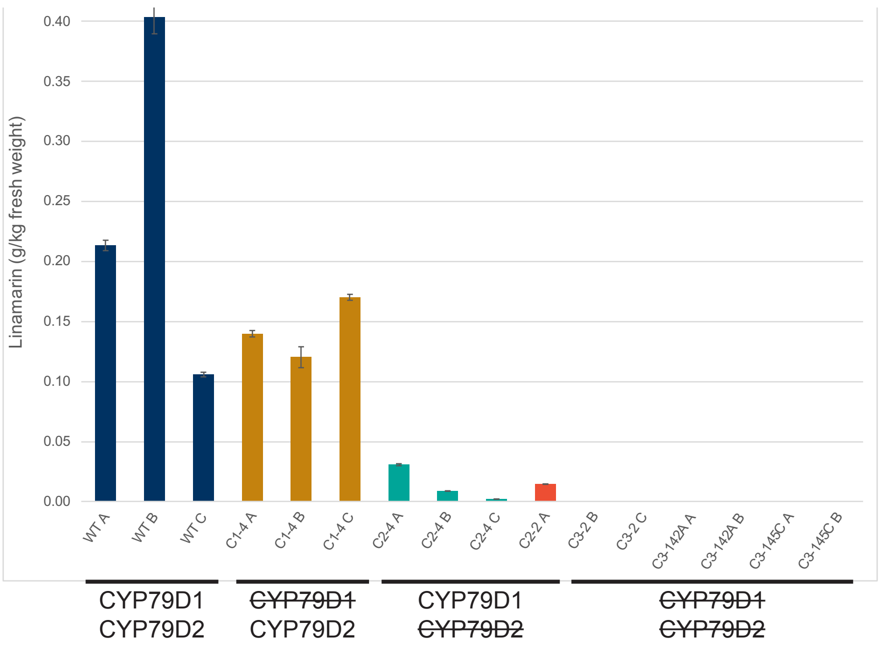


(b)

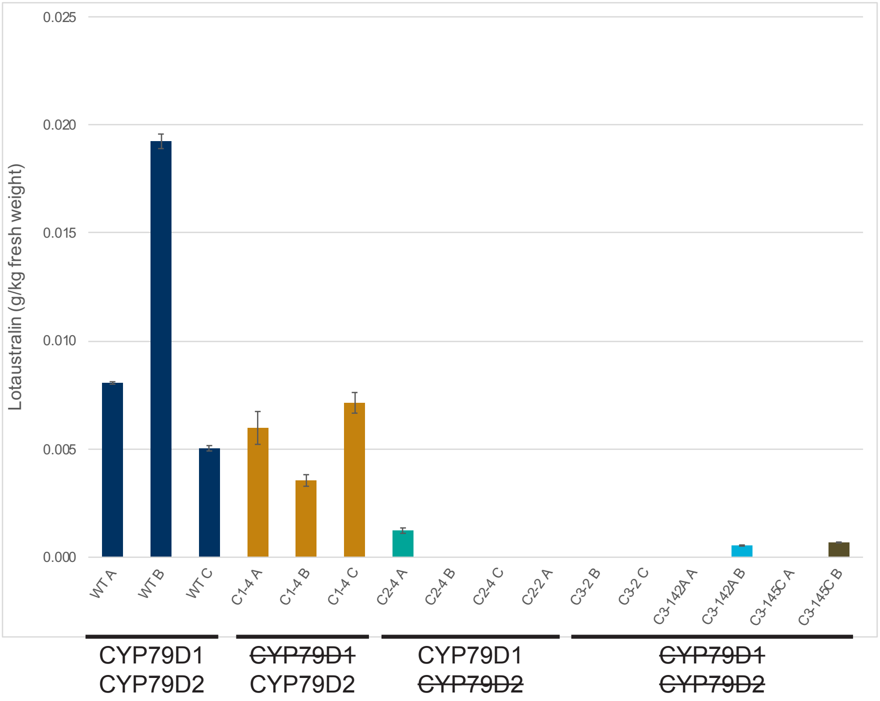


**Figure S5** Cyanogen levels in leaves of edited 60444 *in vitro* plantlets. Linamarin (a) and lotaustralin (b) levels in g/kg fresh weight, as measured via LC-MS. Each bar represents one leaf sample from one plant, as the mean of three technical replicates. Error bars are standard error, calculated from technical replicates. Values below the limit of quantification were treated as 0; standard error was not calculated for these. Line identifiers are followed by a plant identifier (A, B, or C). Bars of the same color are plants of the same line. Edited lines are identified by the CRISPR construct with which they were modified (C1, C2, C3), followed by an index number (e.g. 142A).

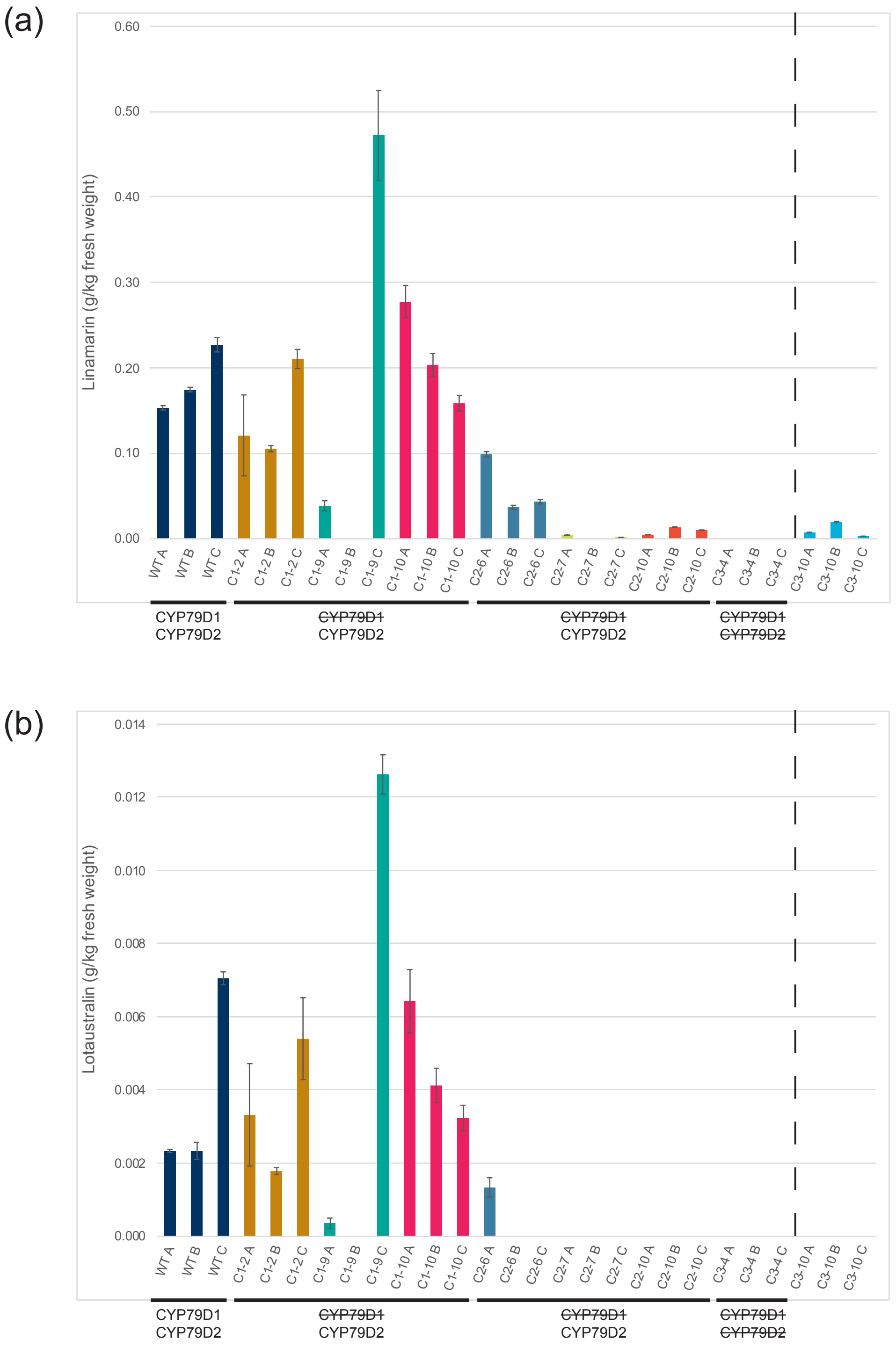


**Figure S6** Cyanogen levels in leaves of edited TME 419 *in vitro* plantlets. Linamarin (a) and lotaustralin (b) levels in g/kg fresh weight, as measured via LC-MS. Each bar represents one leaf sample from one plant, as the mean of three technical replicates. Error bars are standard error, calculated from technical replicates. Values below the limit of quantification were treated as 0; standard error was not calculated for these. Line identifiers are followed by a plant identifier (A, B, or C). Bars of the same color are plants of the same line. Line C3-10 is mosaic for a wildtype *CYP79D2* allele. Edited lines are identified by the CRISPR construct with which they were modified (C1, C2, C3), followed by an index number.

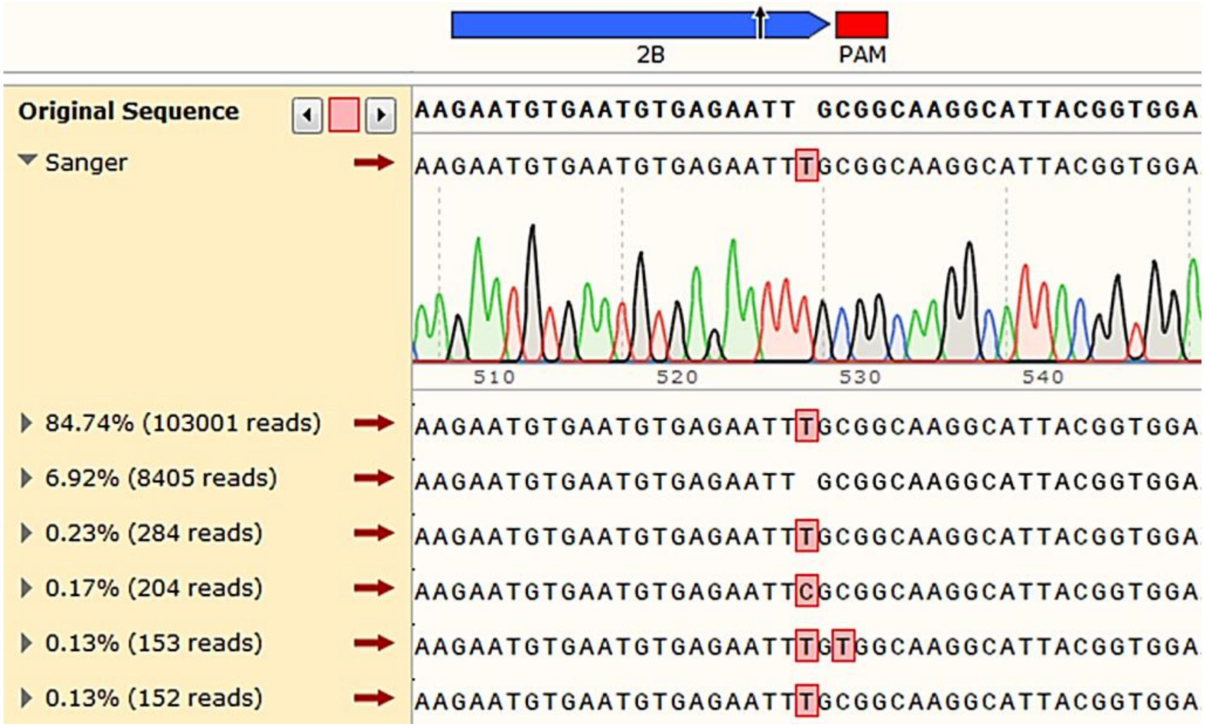


**Figure S7** Sanger and Illumina sequence analysis of TME 419 line C3-10 at target site 2B. Diagram of the protospacer (blue) and protospacer adjacent motif (PAM, red) of the target is aligned to the nucleotide sequence. Black arrow indicates predicted CRISPR-Cas9 cut site. Original sequence (wildtype) is shown in bold letters. Sanger sequence is shown with chromatogram analysis; unique Illumina sequences are shown below. Mismatches highlighted in red boxes. Some distinguishing mismatches between the read sets are outside of the figure window. Map created with SnapGene.

(a) CYP79D1 (b) CYP79D2

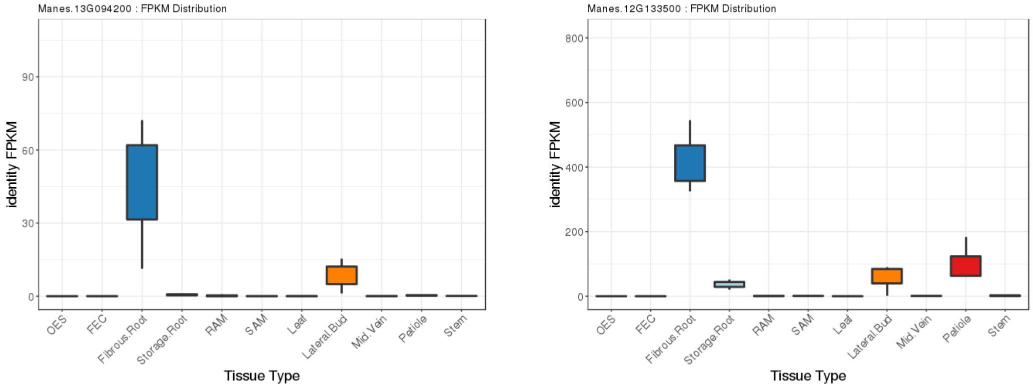


**Figure S8** Tissue-specific transcript expression of *CYP79D1* and *CYP79D2*. Graphs were produced using the Cassava Atlas tool ((Wilson *et al.*, 2017); http://shiny.danforthcenter.org/cassava_atlas). African cassava accession TME 204 was sampled for gene expression 3 months after planting. Each box and whisker plot shows gene expression levels across cassava tissues/organs. FPKM, fragments per kilobase of transcript per million; OES, organized embryogenic structures; FEC, friable embryogenic callus; SAM, shoot apical meristem; RAM, root apical meristem.

### Supplementary Files

**File S1** Genotypes of CRISPR/Cas9-edited 60444 and TME 419 lines. Genotypes at *CYP79D1* and *CYP79D2* target sites identified by category (wildtype, homozygous, bi-allelic, heterozygous, complex) and number of base pairs inserted (i) and/or deleted (d) for each allele. For bi-allelic genotypes with indels of equivalent size but different base pairs, mutations are distinguished by letters “a” and “b”. Deletions that span both target sites are underlined. S (Sanger) or I (Illumina) denotes the type of sequencing applied. Lines originated from the Donald Danforth Plant Science Center (DDPSC) or Innovative Genomics Institute (IGI). WT, wildtype; N.D., not determined.

**File S2** Predicted CYP79D amino acid sequences from selected 60444 and TME 419 mutant lines. Amino acid sequences in FASTA format are based on AM560-2 reference assembly v8.1 (Bredeson *et al.*, 2021). Sequence names are presented in order as accession, gene, line, and, in the case of bi-allelic and heterozygous lines, allele number 1 or 2.

**File S3** LC-MS and picrate readings. LC-MS concentration values reported as <LLOQ (below the lower limit of quantification) were treated as 0 µM. In picrate worksheets, horizontal dotted lines delineate samples of a given line that were analyzed in separate waves of analysis.

**File S4** Python notebook for figures 2–4. Code used for generating cyanide content box and whisker plots, and one-to-one group rank sum comparisons. All data used to generate the plots were gathered from picrate paper assays.
