## Supplementary material for "CRISPR-Cas9-mediated knockout of *CYP79D1* and *CYP79D2* in cassava attenuates toxic cyanogen production": File S4: Python notebook for figs 2-4: Gomezetal_FileS4.ipynb

```
# %% [code]
import io
import numpy as np
import pandas as pd
import seaborn as sns
import matplotlib as mpl
import matplotlib.pyplot as plt
from scipy.stats import ranksums 

# %% [code]
from google.colab import drive
drive.mount('/content/drive')

# %% [markdown]
FOR 60444 PICRATE PLOTS

# %% [code]
# Regular picrate with PPM tsv
uploaded1 = '/content/drive/My Drive/Picrate Assay tsv files/Picrate_assay1_60444.tsv'

# Height (unsused)
uploaded2 = '/content/drive/My Drive/Picrate Assay tsv files/Picrate_assay1_60444_Height.tsv'

# %% [code]
# Store csv as Pandas Dataframe
picrate = pd.read_csv(uploaded1, sep='\t', header=0)
picrate_height = pd.read_csv(uploaded2, sep='\t', header=0)

# %% [code]
picrate_leaf = picrate.loc[picrate["Tissue"] == "leaf"]
picrate_root = picrate.loc[picrate["Tissue"] == "root"]

# %% [code]
picrate_leaf.head()

# %% [code]
picrate_root.head()

# %% [code]
# Dropping DC1-2 + making leaf and root specific df
picrate_root = picrate_root[picrate_root['60444 Genotype'] != 'DC1-2']
picrate_leaf = picrate_leaf[picrate_leaf['60444 Genotype'] != 'DC1-2']
picrate_height = picrate_height[picrate_height['Genotype'] != 'DC1-2']

# %% [code]
# Set sns Theme
sns.set_theme()
rc={'lines.linewidth': 1, 'axes.labelsize': 13, 'axes.titlesize': 15}
sns.set(rc=rc)
my_pal = {"Wildtype": "#00B0DA",  
          "DC1-4": "#FDB515", 
          "DC1-6": "#FDB515", 
          "DC2-4": "#00A598", 
          "DC2-2": "#00A598", 
          "DC3-2": "#ED4E33", 
          "IC3-142A": "#ED4E33",
          "IC3-145C": "#ED4E33"}
# "DC1-2": "#FDB515",

# %% [code]
# Bee Swarm and Box Plots for 60444 Leaf and Root Cyanide Content

sns.set_style("whitegrid")

fig2_3_leaf, ax2_3_leaf = plt.subplots(figsize=(10, 6))
ax2_3_leaf = sns.boxplot(x="60444 Genotype", y="PPM spec direct", data=picrate_leaf, palette=my_pal)
ax2_3_leaf = sns.swarmplot(x='60444 Genotype', y='PPM spec direct', data=picrate_leaf, color="black")
ax2_3_leaf.set_title('60444 Leaf Cyanide Content')
ax2_3_leaf.set_ylabel('Cyanide PPM Spec')
ax2_3_leaf.set_xticklabels(ax2_3_leaf.get_xticklabels(), rotation=45)

plt.savefig('_60444_Leaf.png', dpi=1440)
# plt.savefig('_60444_Leaf.pdf')
plt.show()

fig2_3_root, ax2_3_root = plt.subplots(figsize=(10, 6))
ax2_3_root = sns.boxplot(x="60444 Genotype", y="PPM spec direct", data=picrate_root, palette=my_pal)
ax2_3_root = sns.swarmplot(x='60444 Genotype', y='PPM spec direct', data=picrate_root, color="black")
ax2_3_root.set_title('60444 Root Cyanide Content')
ax2_3_root.set_ylabel('Cyanide PPM Spec')
ax2_3_root.set_xticklabels(ax2_3_root.get_xticklabels(), rotation=45)

plt.savefig('_60444_Root.png', dpi=1440)
# plt.savefig('_60444_Root.pdf')
plt.show()

!mv _60444_Leaf.png _60444_Root.png /content/drive/My\ Drive/
!ls

# %% [markdown]
FOR TME419 PICRATE PLOTS

# %% [code]
# Regular picrate with PPM tsv
uploaded1 = '/content/drive/My Drive/Picrate Assay tsv files/Picrate_assay2_TME419.tsv'

# Height tsv (unused)
uploaded2 = '/content/drive/My Drive/Picrate Assay tsv files/Picrate_assay2_TME419_Height.tsv'

# %% [code]
# Store csv as Pandas Dataframe
picrate = pd.read_csv(uploaded1, sep='\t', header=0)
picrate_height = pd.read_csv(uploaded2, sep='\t', header=0)

# Removing DC1-10 and DC3-2
picrate_height = picrate_height.loc[picrate_height["Genotype"] != "DC1-10"]
picrate_height = picrate_height.loc[picrate_height["Genotype"] != "DC3-2"]

# %% [code]
# Removing DC1-10 and DC3-2 + making leaf and root specific df
picrate_leaf = picrate.loc[picrate["Tissue"] == "leaf"]
picrate_leaf = picrate_leaf.loc[picrate["TME419 Genotype"] != "DC1-10"]
picrate_leaf = picrate_leaf.loc[picrate_leaf["TME419 Genotype"] != "DC3-10"]

picrate_root = picrate.loc[picrate["Tissue"] == "root"]
picrate_root = picrate_root.loc[picrate["TME419 Genotype"] != "DC1-10"]
picrate_root = picrate_root.loc[picrate["TME419 Genotype"] != "DC3-10"]

# %% [code]
picrate_leaf.head()

# %% [code]
picrate_root.head()

# %% [code]
# Set sns Theme
sns.set_theme()
rc={'lines.linewidth': 1, 'axes.labelsize': 13, 'axes.titlesize': 15}
sns.set(rc=rc)

# Set custom palette
my_pal = {"Wildtype": "#00B0DA", 
          "DC1-2": "#FDB515", 
          "DC1-9": "#FDB515", 
          "DC2-6": "#00A598", 
          "DC2-7": "#00A598", 
          "DC2-10": "#00A598", 
          "DC3-4": "#ED4E33"}

# %% [code]
# Bee Swarm and Box Plots for 60444 Leaf and Root Cyanide Content
from google.colab import files
sns.set_style("whitegrid")

fig2_3_leaf, ax2_3_leaf = plt.subplots(figsize=(10, 6))
ax2_3_leaf = sns.boxplot(x="TME419 Genotype", y="ppm spec", data=picrate_leaf, palette=my_pal)
ax2_3_leaf = sns.swarmplot(x='TME419 Genotype', y='ppm spec', data=picrate_leaf, color="black")
ax2_3_leaf.set_title('TME419 Leaf Cyanide Content')
ax2_3_leaf.set_ylabel('Cyanide PPM Spec')
ax2_3_leaf.set_xticklabels(ax2_3_leaf.get_xticklabels(), rotation=45)

fig2_3_leaf.savefig('TME419_Leaf_No_DC3-10.png', dpi=1440)
# fig2_3_leaf.savefig('TME419_Leaf_No_DC3-10.pdf')
plt.show()

fig2_3_root, ax2_3_root = plt.subplots(figsize=(10, 6))
ax2_3_root = sns.boxplot(x="TME419 Genotype", y='ppm spec', data=picrate_root, palette=my_pal)
ax2_3_root = sns.swarmplot(x='TME419 Genotype', y='ppm spec', data=picrate_root, color="black")
ax2_3_root.set_title('TME419 Root Cyanide Content')
ax2_3_root.set_ylabel('Cyanide PPM Spec')
ax2_3_root.set_xticklabels(ax2_3_root.get_xticklabels(), rotation=45)

fig2_3_root.savefig('TME419_Root_No_DC3-10.png', dpi=1440)
# fig2_3_root.savefig('TME419_Root_No_DC3-10.pdf')
plt.show()

!mv TME419_Leaf_No_DC3-10.png TME419_Root_No_DC3-10.png /content/drive/My\ Drive/
!ls

# %% [markdown]
FOR SURVEY 5-2-21 PICRATE PLOTS

# %% [code]
# Regular picrate with PPM tsv
uploaded1 = '/content/drive/My Drive/Picrate Assay tsv files/Picrate_assay_survey_5_21_2021.tsv'

# Height tsv (unused)
uploaded2 = '/content/drive/My Drive/Picrate Assay tsv files/Picrate_assay_survey_HEIGHT_5_21_2021.tsv'

# %% [code]
# Store csv as Pandas Dataframe
# Removing Oko-iyawo and Kibandameno
picrate = pd.read_csv(uploaded1, sep='\t', header=0)
picrate_height = pd.read_csv(uploaded2, sep='\t', header=0)
picrate_height = picrate_height.loc[picrate_height["Genotype "] != "Oko-iyawo"]
picrate_height = picrate_height.loc[picrate_height["Genotype "] != "Kibandameno"]

# %% [code]
# Dropping Oko-iyawo and Kibandameno + making leaf and root specific df
picrate_leaf = picrate.loc[picrate["Tissue"] == "leaf"]
picrate_leaf = picrate_leaf.loc[picrate_leaf["Genotype"] != "Oko-iyawo"]
picrate_leaf = picrate_leaf.loc[picrate_leaf["Genotype"] != "Kibandameno"]

picrate_root = picrate.loc[picrate["Tissue"] == "root"]
picrate_root = picrate_root.loc[picrate_root["Genotype"] != "Oko-iyawo"]
picrate_root = picrate_root.loc[picrate_root["Genotype"] != "Kibandameno"]

# %% [code]
picrate_leaf.head()

# %% [code]
picrate_root.head()

# %% [code]
# Set sns Theme
sns.set_theme()
rc={'lines.linewidth': 1, 'axes.labelsize': 13, 'axes.titlesize': 15}
sns.set(rc=rc)

# Set custom palette
my_pal = {"60444": "#00B0DA", 
          "TME419": "#FDB515", 
          "TMS91/02324": "#00A598", 
          "TME3": "#ED4E33", 
          "TMS98/0505": "#c9581b",
          "MCol22": "#c9581b",
          "MCol2215": "#c9581b",
          }

# %% [code]
# Bee Swarm and Box Plots for 60444 Leaf and Root Cyanide Content
from google.colab import files
sns.set_style("whitegrid")

fig2_3_leaf, ax2_3_leaf = plt.subplots(figsize=(10, 6))
ax2_3_leaf = sns.boxplot(x="Genotype", y="ppm spec", data=picrate_leaf, palette='colorblind')
ax2_3_leaf = sns.swarmplot(x='Genotype', y='ppm spec', data=picrate_leaf, color="black")
ax2_3_leaf.set_title('Leaf Cyanide Content')
ax2_3_leaf.set_ylabel('Cyanide PPM Spec')
ax2_3_leaf.set_xticklabels(ax2_3_leaf.get_xticklabels(), rotation=45, fontsize=8)

fig2_3_leaf.savefig('Survey_5_2_21_Leaf.png', dpi=1440)
# fig2_3_leaf.savefig('Survey_5_2_21_Leaf.pdf')
plt.show()

fig2_3_root, ax2_3_root = plt.subplots(figsize=(10, 6))
ax2_3_root = sns.boxplot(x="Genotype", y='ppm spec', data=picrate_root, palette='colorblind')
ax2_3_root = sns.swarmplot(x='Genotype', y='ppm spec', data=picrate_root, color="black")
ax2_3_root.set_title('Root Cyanide Content')
ax2_3_root.set_ylabel('Cyanide PPM Spec')
ax2_3_root.set_xticklabels(ax2_3_root.get_xticklabels(), rotation=45, fontsize=8)

fig2_3_root.savefig('Survey_5_2_21_Root.png', dpi=1440)
# fig2_3_root.savefig('Survey_5_2_21_Root.pdf')
plt.show()

!mv Survey_5_2_21_Leaf.png Survey_5_2_21_Root.png /content/drive/My\ Drive/
!ls

# %% [markdown]
FOR WILCOX RANK SUM PLOTS (ALL PLOTS)

# %% [code]
# import data
gen_60444 = '/content/drive/My Drive/Picrate Assay tsv files/Picrate_assay1_60444.tsv'
gen_TME419 = '/content/drive/My Drive/Picrate Assay tsv files/Picrate_assay2_TME419.tsv'
gen_survey = '/content/drive/My Drive/Picrate Assay tsv files/Picrate_assay3.tsv'
gen_survey_5_2 = '/content/drive/My Drive/Picrate Assay tsv files/Picrate_assay_survey_5_21_2021.tsv'

# convert to pandas dataframe
gen_60444_df = pd.read_csv(gen_60444, sep='\t', header=0)
gen_TME419_df = pd.read_csv(gen_TME419, sep='\t', header=0)
gen_survey_df = pd.read_csv(gen_survey, sep='\t', header=0)
gen_survey_5_2_df = pd.read_csv(gen_survey_5_2, sep='\t', header=0)

# grab only the root data
gen_60444_root_df = gen_60444_df.loc[gen_60444_df["Tissue"] == "root"]
gen_TME419_root_df = gen_TME419_df.loc[gen_TME419_df["Tissue"] == "root"]
gen_survey_root_df = gen_survey_df.loc[gen_survey_df["Tissue"] == "root"]
gen_survey_5_2_root_df = gen_survey_5_2_df.loc[gen_survey_5_2_df["Tissue"] == "root"]


# %% [code]
# turn all groups of data into arrays 
# 60444
WT_60444_root = gen_60444_root_df[gen_60444_root_df['60444 Genotype'] == 'Wildtype']['PPM spec direct'].dropna()
C1_4_60444_root = gen_60444_root_df[gen_60444_root_df['60444 Genotype'] == 'DC1-4']['PPM spec direct'].dropna()
C1_6_60444_root = gen_60444_root_df[gen_60444_root_df['60444 Genotype'] == 'DC1-6']['PPM spec direct'].dropna()
C2_2_60444_root = gen_60444_root_df[gen_60444_root_df['60444 Genotype'] == 'DC2-2']['PPM spec direct'].dropna()
C2_4_60444_root = gen_60444_root_df[gen_60444_root_df['60444 Genotype'] == 'DC2-4']['PPM spec direct'].dropna()
C3_2_60444_root = gen_60444_root_df[gen_60444_root_df['60444 Genotype'] == 'DC3-2']['PPM spec direct'].dropna()
C3_142A_60444_root = gen_60444_root_df[gen_60444_root_df['60444 Genotype'] == 'IC3-142A']['PPM spec direct'].dropna()
C3_145C_60444_root = gen_60444_root_df[gen_60444_root_df['60444 Genotype'] == 'IC3-145C']['PPM spec direct'].dropna()

sample_arr_60444 = [WT_60444_root, C1_4_60444_root, C1_6_60444_root, C2_2_60444_root, C2_4_60444_root,
                    C3_2_60444_root, C3_142A_60444_root, C3_145C_60444_root]

# TME419
WT_TME419_root = gen_TME419_root_df[gen_TME419_root_df['TME419 Genotype'] == 'Wildtype']['ppm spec'].dropna()
C1_2_TME419_root = gen_TME419_root_df[gen_TME419_root_df['TME419 Genotype'] == 'DC1-2']['ppm spec'].dropna()
C1_9_TME419_root = gen_TME419_root_df[gen_TME419_root_df['TME419 Genotype'] == 'DC1-9']['ppm spec'].dropna()
C2_6_TME419_root = gen_TME419_root_df[gen_TME419_root_df['TME419 Genotype'] == 'DC2-6']['ppm spec'].dropna()
C2_7_TME419_root = gen_TME419_root_df[gen_TME419_root_df['TME419 Genotype'] == 'DC2-7']['ppm spec'].dropna()
C2_10_TME419_root = gen_TME419_root_df[gen_TME419_root_df['TME419 Genotype'] == 'DC2-10']['ppm spec'].dropna()
C3_4_TME419_root = gen_TME419_root_df[gen_TME419_root_df['TME419 Genotype'] == 'DC3-4']['ppm spec'].dropna()
# C3_10_TME419_root = gen_TME419_root_df[gen_TME419_root_df['TME419 Genotype'] == 'DC3-10']['ppm spec'].dropna()

sample_arr_TME419 = [WT_TME419_root, C1_2_TME419_root, C1_9_TME419_root, C2_6_TME419_root, C2_7_TME419_root, 
                    C2_10_TME419_root, C3_4_TME419_root]  

# Survey
GENOTYPE_60444_root = gen_survey_root_df[gen_survey_root_df['Accession'] == '60444']['ppm spec'].dropna()
TME419_root = gen_survey_root_df[gen_survey_root_df['Accession'] == 'TME419']['ppm spec'].dropna()
TMS_91_02324_root = gen_survey_root_df[gen_survey_root_df['Accession'] == 'TMS 91/02324']['ppm spec'].dropna()
TME3_root = gen_survey_root_df[gen_survey_root_df['Accession'] == 'TME3']['ppm spec'].dropna()
Oko_iyawo_root = gen_survey_root_df[gen_survey_root_df['Accession'] == 'Oko-iyawo']['ppm spec'].dropna()
TMS_98_0505_root = gen_survey_root_df[gen_survey_root_df['Accession'] == 'TMS 98/0505']['ppm spec'].dropna() 

sample_arr_survey = [GENOTYPE_60444_root, TME419_root, TMS_91_02324_root, TME3_root, Oko_iyawo_root, 
                    TMS_98_0505_root]

# Survey 5-2-2021
S_5_2_60444_root = gen_survey_5_2_root_df[gen_survey_5_2_root_df['Genotype'] == '60444']['ppm spec'].dropna()
S_5_2_TME419_root = gen_survey_5_2_root_df[gen_survey_5_2_root_df['Genotype'] == 'TME419']['ppm spec'].dropna()
S_5_2_TMS_91_02324_root = gen_survey_5_2_root_df[gen_survey_5_2_root_df['Genotype'] == 'TMS91/02324']['ppm spec'].dropna()
S_5_2_TME3_root = gen_survey_5_2_root_df[gen_survey_5_2_root_df['Genotype'] == 'TME3']['ppm spec'].dropna()
S_5_2_TMS_98_0505_root = gen_survey_5_2_root_df[gen_survey_5_2_root_df['Genotype'] == 'TMS98/0505']['ppm spec'].dropna()
S_5_2_MCol22_root = gen_survey_5_2_root_df[gen_survey_5_2_root_df['Genotype'] == 'MCol22']['ppm spec'].dropna()
S_5_2_MCol2215_root = gen_survey_5_2_root_df[gen_survey_5_2_root_df['Genotype'] == 'MCol2215']['ppm spec'].dropna()

sample_arr_survey_5_2 = [S_5_2_60444_root, S_5_2_TME419_root, S_5_2_TMS_91_02324_root, S_5_2_TME3_root, S_5_2_TMS_98_0505_root, 
                    S_5_2_MCol22_root, S_5_2_MCol2215_root]


# %% [code]
# function for Wilcoxon rank-sum test
def rank_sum_test_all(sample_array):
  out_matrix_p_values = []

  for i in range(len(sample_array)):
    out_row_in_matrix = []
    for j in range(len(sample_array)):
      p_value = ranksums(sample_array[i], sample_array[j])
      out_row_in_matrix.append(p_value[1])
    out_matrix_p_values.append(out_row_in_matrix)

  return out_matrix_p_values


# %% [code]
cmap1 = mpl.colors.ListedColormap(['#aa1016'])
cmap2 = mpl.colors.ListedColormap(['#d52221'])
cmap3 = mpl.colors.ListedColormap(['#f44f39'])
cmap4 = mpl.colors.ListedColormap(['#fc8161'])
cmap5 = mpl.colors.ListedColormap(['#fcaf93'])
cmap6 = mpl.colors.ListedColormap(['#fedbcc'])
cmap7 = mpl.colors.ListedColormap(['#dbe9f6'])
cmap8 = mpl.colors.ListedColormap(['#bad6eb'])
cmap9 = mpl.colors.ListedColormap(['#89bedc'])
cmap10 = mpl.colors.ListedColormap(['#539ecd'])
cmap11 = mpl.colors.ListedColormap(['#2b7bba'])
cmap12 = mpl.colors.ListedColormap(['#0b559f'])

# %% [code]
# getting the data 60444
p_values = rank_sum_test_all(sample_arr_60444)

# mask = np.zeros_like(p_values)
# mask[np.triu_indices_from(mask)] = True
fig, ax = plt.subplots(figsize=(10,10)) 


sns.heatmap(p_values, mask=np.array(p_values) < 0.0001, cmap=cmap12, cbar=False, annot=True)
sns.heatmap(p_values, mask=np.array(p_values) < 0.0005, cmap=cmap11, cbar=False)
sns.heatmap(p_values, mask=np.array(p_values) < 0.001, cmap=cmap10, cbar=False)
sns.heatmap(p_values, mask=np.array(p_values) < 0.005, cmap=cmap9, cbar=False)
sns.heatmap(p_values, mask=np.array(p_values) < 0.01, cmap=cmap8, cbar=False, annot=True)
sns.heatmap(p_values, mask=np.array(p_values) < 0.03, cmap=cmap7, cbar=False, annot=True)

sns.heatmap(p_values, mask=np.array(p_values) < 0.05, cmap=cmap6, cbar=False)
sns.heatmap(p_values, mask=np.array(p_values) < 0.1, cmap=cmap5, cbar=False)
sns.heatmap(p_values, mask=np.array(p_values) < 0.3, cmap=cmap4, cbar=False)
sns.heatmap(p_values, mask=np.array(p_values) < 0.5, cmap=cmap3, cbar=False)
sns.heatmap(p_values, mask=np.array(p_values) < 0.7, cmap=cmap2, cbar=False)
sns.heatmap(p_values, mask=np.array(p_values) < 0.9, cmap=cmap1, cbar=False, annot=True)

plt.title("Wilcoxon rank-sum p-value between 60444 genotypes")
plt.xlabel('60444 genotypes (WT = 0)')
plt.ylabel('60444 genotypes (WT = 0)')

plt.savefig('_60444_Root_W_ranksum.png', dpi=1440)
# plt.savefig('_60444_Root_W_ranksum.pdf')
plt.show()

!mv _60444_Root_W_ranksum.png /content/drive/My\ Drive/
!ls

# %% [code]
# getting the data TME419
p_values = rank_sum_test_all(sample_arr_TME419)

# mask = np.zeros_like(p_values)
# mask[np.triu_indices_from(mask)] = True
fig, ax = plt.subplots(figsize=(10,10)) 

sns.heatmap(p_values, mask=np.array(p_values) < 0.0001, cmap=cmap12, cbar=False, annot=True)
sns.heatmap(p_values, mask=np.array(p_values) < 0.0005, cmap=cmap11, cbar=False)
sns.heatmap(p_values, mask=np.array(p_values) < 0.001, cmap=cmap10, cbar=False)
sns.heatmap(p_values, mask=np.array(p_values) < 0.005, cmap=cmap9, cbar=False)
sns.heatmap(p_values, mask=np.array(p_values) < 0.01, cmap=cmap8, cbar=False, annot=True)
sns.heatmap(p_values, mask=np.array(p_values) < 0.03, cmap=cmap7, cbar=False, annot=True)

sns.heatmap(p_values, mask=np.array(p_values) < 0.05, cmap=cmap6, cbar=False)
sns.heatmap(p_values, mask=np.array(p_values) < 0.1, cmap=cmap5, cbar=False)
sns.heatmap(p_values, mask=np.array(p_values) < 0.3, cmap=cmap4, cbar=False)
sns.heatmap(p_values, mask=np.array(p_values) < 0.5, cmap=cmap3, cbar=False)
sns.heatmap(p_values, mask=np.array(p_values) < 0.7, cmap=cmap2, cbar=False)
sns.heatmap(p_values, mask=np.array(p_values) < 0.9, cmap=cmap1, cbar=False, annot=True)

plt.title("Wilcoxon rank-sum p-value between TME419 genotypes")
plt.xlabel('TME419 genotypes (WT = 0)')
plt.ylabel('TME419 genotypes (WT = 0)')

plt.savefig('TME419_Root_W_ranksum_No_DC3-10.png', dpi=1440)
# plt.savefig('TME419_Root_W_ranksum_No_DC3-10.pdf')
plt.show()

!mv TME419_Root_W_ranksum_No_DC3-10.png /content/drive/My\ Drive/
!ls

# %% [code]
# # getting the data survey plants
# p_values = rank_sum_test_all(sample_arr_survey)

# # mask = np.zeros_like(p_values)
# # mask[np.triu_indices_from(mask)] = True
# fig, ax = plt.subplots(figsize=(10,10)) 

# sns.heatmap(p_values, mask=np.array(p_values) < 0.0001, cmap=cmap12, cbar=False, annot=True)
# sns.heatmap(p_values, mask=np.array(p_values) < 0.0005, cmap=cmap11, cbar=False)
# sns.heatmap(p_values, mask=np.array(p_values) < 0.001, cmap=cmap10, cbar=False)
# sns.heatmap(p_values, mask=np.array(p_values) < 0.005, cmap=cmap9, cbar=False)
# sns.heatmap(p_values, mask=np.array(p_values) < 0.01, cmap=cmap8, cbar=False, annot=True)
# sns.heatmap(p_values, mask=np.array(p_values) < 0.03, cmap=cmap7, cbar=False, annot=True)

# sns.heatmap(p_values, mask=np.array(p_values) < 0.05, cmap=cmap6, cbar=False)
# sns.heatmap(p_values, mask=np.array(p_values) < 0.1, cmap=cmap5, cbar=False)
# sns.heatmap(p_values, mask=np.array(p_values) < 0.3, cmap=cmap4, cbar=False)
# sns.heatmap(p_values, mask=np.array(p_values) < 0.5, cmap=cmap3, cbar=False)
# sns.heatmap(p_values, mask=np.array(p_values) < 0.7, cmap=cmap2, cbar=False)
# sns.heatmap(p_values, mask=np.array(p_values) < 0.9, cmap=cmap1, cbar=False, annot=True)

# plt.title("Wilcoxon rank-sum p-value between survey genotypes")
# plt.xlabel('Genotypes (60444 = 0)')
# plt.ylabel('Genotypes (60444 = 0)')

# plt.savefig('Survey_Root_W_ranksum.png', dpi=1080)
# plt.show()

# !mv Survey_Root_W_ranksum.png /content/drive/My\ Drive/
# !ls

# %% [code]
# getting the data survey plants
p_values = rank_sum_test_all(sample_arr_survey_5_2)

# mask = np.zeros_like(p_values)
# mask[np.triu_indices_from(mask)] = True
fig, ax = plt.subplots(figsize=(10,10)) 

sns.heatmap(p_values, mask=np.array(p_values) < 0.0001, cmap=cmap12, cbar=False, annot=True)
sns.heatmap(p_values, mask=np.array(p_values) < 0.0005, cmap=cmap11, cbar=False)
sns.heatmap(p_values, mask=np.array(p_values) < 0.001, cmap=cmap10, cbar=False)
sns.heatmap(p_values, mask=np.array(p_values) < 0.005, cmap=cmap9, cbar=False)
sns.heatmap(p_values, mask=np.array(p_values) < 0.01, cmap=cmap8, cbar=False, annot=True)
sns.heatmap(p_values, mask=np.array(p_values) < 0.03, cmap=cmap7, cbar=False, annot=True)

sns.heatmap(p_values, mask=np.array(p_values) < 0.05, cmap=cmap6, cbar=False)
sns.heatmap(p_values, mask=np.array(p_values) < 0.1, cmap=cmap5, cbar=False)
sns.heatmap(p_values, mask=np.array(p_values) < 0.3, cmap=cmap4, cbar=False)
sns.heatmap(p_values, mask=np.array(p_values) < 0.5, cmap=cmap3, cbar=False)
sns.heatmap(p_values, mask=np.array(p_values) < 0.7, cmap=cmap2, cbar=False)
sns.heatmap(p_values, mask=np.array(p_values) < 0.9, cmap=cmap1, cbar=False, annot=True)

plt.title("Wilcoxon rank-sum p-value between survey genotypes 5-2-2021")
plt.xlabel('Genotypes (60444 = 0)')
plt.ylabel('Genotypes (60444 = 0)')

plt.savefig('Survey_5_2_21_Root_W_ranksum.png', dpi=1440)
# plt.savefig('Survey_5_2_21_Root_W_ranksum.pdf')
plt.show()

!mv Survey_5_2_21_Root_W_ranksum.png /content/drive/My\ Drive/
!ls

# %% [markdown]
$W = min(W_1, W_2) \textrm{, where} \ W_1 \ \textrm{and} \ W_2 \ \textrm{are the rank sums of } \\ \textrm{Group 1 and 2, respectively.} \\ \ \\
\textrm{W is approximated to be normally distributed with mean } \\ 
\mu_w = \frac{n_1 (n_1 + n_2 + 1}{2} \textrm{ and standard deviation } 
\sigma_w = \sqrt{\frac{n_1 n_2 (n_1 + n_2 + 1)}{12}}. \\ \ \\ 
\textrm{Dist } Z_0 \textrm{ is approximatly normal with mean 0 and variance 1.} \\ \ \\
Z_0 = \frac{W - \mu_w}{\sigma_w},
\textrm{ can be calculated and a corresponding p-score} \\
\textrm{ can be found.}$
```
